# Inflammation of the retinal pigment epithelium drives early-onset photoreceptor degeneration in *Mertk*-associated retinitis pigmentosa

**DOI:** 10.1101/2022.10.08.511415

**Authors:** Maria E. Mercau, Yemsratch T. Akalu, Francesca Mazzoni, Gavin Gyimesi, Emily Alberto, Yong Kong, Brian P. Hafler, Silvia C. Finnemann, Carla V. Rothlin, Sourav Ghosh

## Abstract

Severe, early-onset photoreceptor (PR) degeneration associated with *MERTK* mutations is thought to result from failed phagocytosis by retinal pigment epithelium (RPE). Notwithstanding, the severity and onset of PR degeneration in mouse models of *Mertk* ablation is determined by the hypomorphic expression or the loss of the *Mertk* paralog *Tyro3*. Here we find that loss of *Mertk* and reduced expression/loss of *Tyro3* led to RPE inflammation even before eye-opening. Incipient RPE inflammation cascaded to involve microglia activation and PR degeneration with monocyte infiltration. Inhibition of RPE inflammation with the JAK1/2 inhibitor ruxolitinib mitigated PR degeneration in *Mertk*^-/-^ mice. Neither inflammation nor severe, early-onset PR degeneration were observed in mice with defective phagocytosis alone. Thus, inflammation drives severe, early-onset PR degeneration-associated with *Mertk* loss of function.

## Introduction

Retinitis pigmentosa (RP) is a group of heterogenous degenerative diseases that lead to rod and cone photoreceptor (PR) dystrophy, night blindness, progressive visual field constriction and ultimately central vision loss ^1^. RP is known to be primarily a hereditary disorder with over 45 different genes implicated ^2^. One such gene is *MERTK* ^3^. RP associated with *MERTK* mutations is usually severe and has been observed in patients as early as 3 years of age ^3-6^. Macular involvement is rapid ^3,4^. Severe, early-onset retinal degeneration is also observed in a laboratory model of spontaneous inherited retinal degeneration, the Royal College of Surgeons’ (RCS) rat ^7^. RCS rats develop normal vision with no detectable differences in electroretinogram (ERG) at postnatal day (P) 15 ^7^. ERG changes are detected in RCS rats by P18 and animals are completely blind by P60 ^7^. *In vitro* phagocytosis assays using retinal pigment epithelial (RPE) cells from RCS rats revealed defective engulfment of PR outer segment fragments (POS) in comparison to RPE cells derived from wild type (WT) rats ^8^. It was discovered that the RCS rat carries a ∼1.5 kB deletion that includes the *Mertk* locus ^9,10^ and that MERTK was required for RPE phagocytosis ^11^. The genetic targeting and ablation of *Mertk* in 129P2/OLAHsd (129) embryonic stem (ES) cell-derived mice (henceforth referred to as *Mertk* ^-/-V1^ mice) results in a similar, severe, early-onset retinal degeneration ^12^. ERG recordings revealed that *Mertk* ^-/-V1^ mice displayed early signs of impairment in retinal function at P20 when compared to WT mice ^12^. Based on these observations, it is postulated that the failure of POS engulfment and accumulation of POS debris due to the absence of *Mertk* result in PR degeneration in *Mertk* ^-/-V1^ mice. Nonetheless, not all *MERTK* mutations have a severe, early-onset phenotype. For example, Tschernutter *et al*. identified a family with a milder form of *MERTK* mutation-associated RP ^5^. Additionally, two missense *MERTK* mutations were found in an RP patient but the mother, who had identical mutations, was unaffected ^13^. Such findings raise the possibility that genetic modifiers may determine the severity of RP and time to vision loss.

RPE cells express, in addition to MERTK, other phagocytic receptors such as αvβ5 integrin ^14^, and CD81, which acts as integrin co-receptor ^15^. The MERTK paralog TYRO3 is also expressed in the RPE ^16,17^ and indicated to have a functional role in POS uptake ^17^. Intriguingly, not all mouse models with genetic ablation of phagocytic receptors display the severe, early-onset retinal degeneration phenotype observed in *Mertk* ^-/-V1^ mice. We have previously demonstrated that the *Itgb5* ^-/-^ mouse line exhibits significant RPE phagocytosis defects *in vivo* and *in vitro* ^14,18^. Nonetheless, ERGs from *Itgb5* ^-/-^ and WT mice indicated that both mouse lines had comparable vision at 4 months of age ^14^. *Itgb5* ^-/-^ and WT retina gross morphology was intact even at 12 months of age; however, *Itgb5* ^-/-^ RPE accumulated excessive levels of lipofuscin and ERGs showed loss of PR function with age ^14^. CD81 blockade or silencing inhibited αvβ5 integrin and thus POS uptake by cultured RPE ^15^. However, to date, there are no reports on PR degeneration in *Cd81* ^-/-^ mice. Instead, there was a modest increase in RPE cell and nuclear density and an increase in the number of multinucleated RPE in *Cd81* ^-/-^ eyes ^19,20^. Genetic ablation of the αvβ5 integrin ligand, MFG-E8, yielded the same phagocytic defects as in αvβ5 receptor-deficient mice, but did not show lipofuscin or vision decline by 12 months of age ^18^. Recently, POS phagosome processing by the RPE has been shown to depend on LC3 lipidation and Beclin1, but not on ULK1, ^21^. ERG recordings indicated similar visual function in 3-month old *Vmd2*-cre^+^ *Atg5* ^f/f^ mice and control mice, although retinal function decreased in *Vmd2*-cre^+^ *Atg5* ^f/f^ mice by 4 months of age ^21^. There was no sign of PR degeneration in *Vmd2*-cre^+^ *Atg5* ^f/f^ mice by 6 months of age ^21^. Collectively, these data suggest that PR degeneration resulting from RPE phagocytosis defects is progressive and age-related. By contrast, PR degeneration in the absence of MERTK is severe and early-onset. Therefore, we hypothesized that MERTK may have additional role(s), distinct from its role in phagocytosis. In addition, we proposed that the impact of this function of MERTK on retinal physiology is influenced by modifiers and is crucial for the maintenance of retinal homeostasis.

We and others have identified either the concomitant hypomorphic expression of *Tyro 3* (in *Mertk* ^-/-V1^ mice) or the loss-of-function of *Tyro3* (in *Mertk* ^-/-V2^ *Tyro3* ^-/-V2^ mice) as a modifier of severe, early-onset PR degeneration in mice with the loss of *Mertk* function ^17,22^. Here we show that RPE inflammation is a distinct feature associated with severe, early-onset PR degeneration in *Mertk* ^-/-V1^ and *Mertk* ^-/-V2^ *Tyro3*^-/-V2^ mice. RPE inflammation is detected early, before histological evidence of retinal degeneration, and even before eye-opening. This initial RPE inflammation, subsequently, manifests as microglial activation and ultimately as monocytic infiltration into the retina. Importantly, neither severe, early-onset retinal degeneration, nor exacerbated RPE inflammation is a feature of *Mertk* ^-/-V2^ mice, or of mouse models with genetic ablation of phagocytic receptors such as *Tyro3* ^-/-V1^ or *Itgb5* ^-/-^ mice. The fact that in *Mertk* ^-/-V1^ and *Mertk* ^-/-V2^ *Tyro3* ^-/-V2^ mice RPE inflammation is detected as early as P10, when POS phagocytosis is minimal in rodents, taken together with the lack of exaggerated inflammation in *Mertk* ^-/-V2^, *Tyro3* ^-/-V1^ or *Itgb5* ^-/-^ mice, indicate that this inflammation is not a consequence of failed POS phagocytosis. *Mertk* and *Tyro3* are well-characterized negative regulators of inflammatory cytokine signaling such as type I and II interferons (IFNs), IL-6 and TNF–α. Treatment with ruxolitinib, an FDA-approved JAK1/2 inhibitor, of *Mertk* ^-/-V1^ mice resulted in successful partial preservation of the retinal histology, providing proof-of-concept that inflammation is an important aspect of the etiology of severe, early-onset PR degeneration-associated with loss of function of *Mertk*. In conclusion, our results indicate that *Mertk* and *Tyro3* function redundantly in an anti-inflammatory mechanism in the RPE that is critical for retinal homeostasis. Targeting this inflammatory response in RP with severe, early-onset characteristics might provide a therapeutic avenue to mitigate disease progression.

## Results

### *Mertk* ^-/-V2^ *Tyro3* ^-/-V2^ and *Mertk* ^-/-V1^ mice display severe, early-onset PR degeneration in contrast to progressive peripheral PR degeneration in *Mertk* ^-/-V2^ mice

We previously reported that *Mertk* ^-/-V2^ *Tyro3* ^-/-V2^ mice displayed retinal degeneration ^22^. To define the onset of PR degeneration in *Mertk* ^-/-V2^ *Tyro3* ^-/-V2^ mice we histologically determined the ONL thickness at different postnatal timepoints, by hematoxylin and eosin (H&E) staining (**Fig. 1**). Measurements were taken in areas representing the central, intermediate and peripheral retina (**Fig. 1A**). We observed significant decrease in ONL thickness in *Mertk* ^-/-V2^ *Tyro3* ^-/-V2^ mice as compared to C57BL/6 (B6) WT mice at P35 (**Fig. 1B**). The decrease in ONL thickness in *Mertk* ^-/-V2^ *Tyro3* ^-/-V2^ mice was similar to that observed in *Mertk* ^-/-V1^ mice at P35. This decrease in ONL thickness in *Mertk* ^-/-V2^ *Tyro3* ^-/-V2^ mice, as compared to B6 WT mice, was also observed at P30 and was replicated in *Mertk* ^-/-V1^ mice at this age (**Fig. 1B**). However, ONL thickness at P25 was indistinguishable between B6 WT, *Mertk*^-/-V2^ *Tyro3* ^-/-V2^ and *Mertk* ^-/-V1^ mice (**Fig. 1B**). These results indicate that the severity and onset of retinal degeneration in *Mertk* ^-/-V2^ *Tyro3* ^-/-V2^ and *Mertk* ^-/-V1^ mice were identical across the retina. Therefore, the loss of *Mertk* in the presence of the modifier hypomorphic expression or loss-of-function of *Tyro3* results in identical severe, early-onset PR degeneration.

**Fig. 1.**
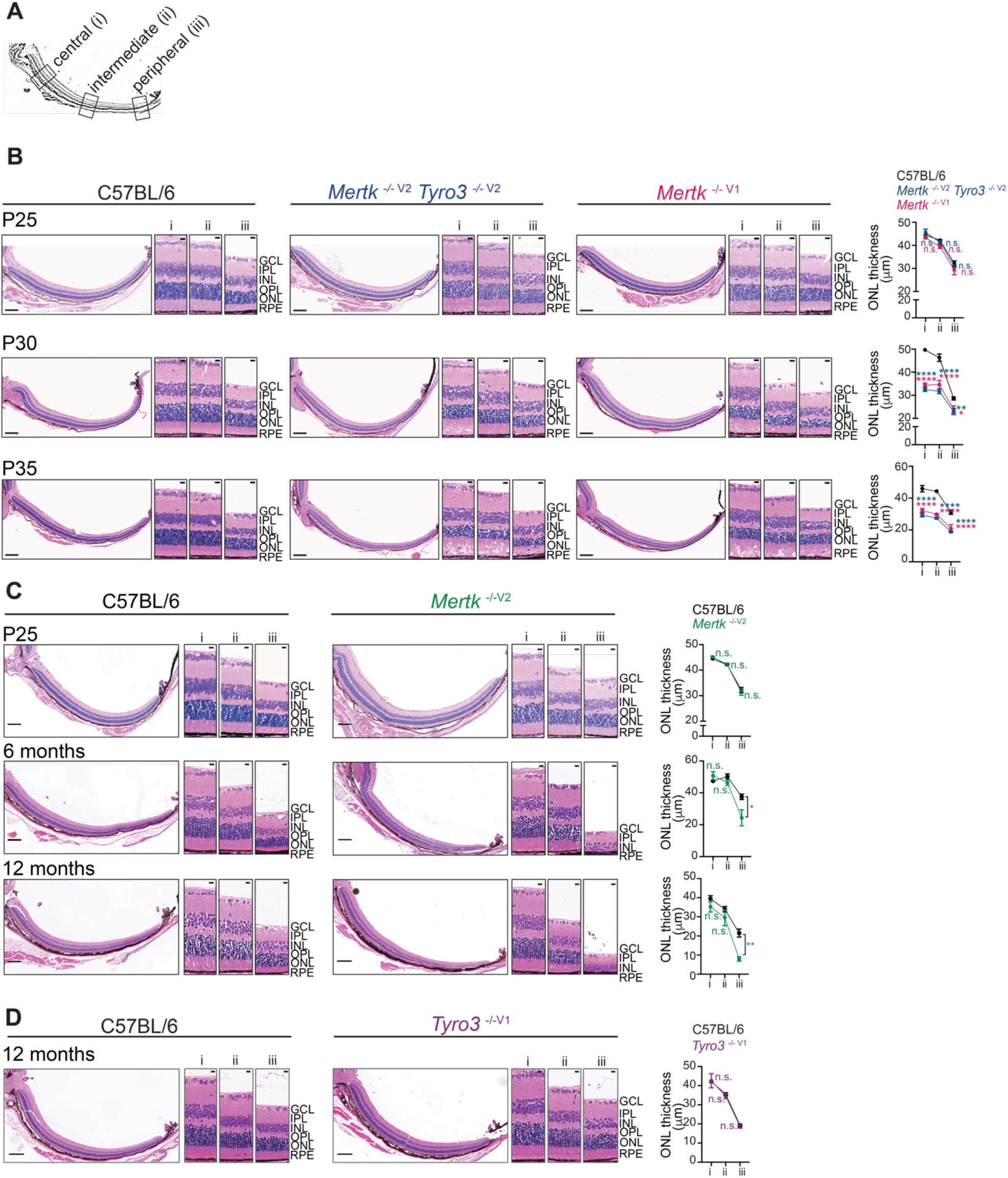
Loss of *Mertk* and concomitant loss or hypomorphic expression of *Tyro3* leads to severe, early-onset PR degeneration. **(A)** Schematic of eye transversal sections indicating the regions of the central (i), intermediate (ii) and peripheral (iii) retina shown and quantified for ONL thickness. **(B)** Eye transversal sections from postnatal day (P) 25, P30, and P35 C57BL/6 WT, *Mertk* ^-/-V2^ *Tyro3* ^-/-V2^ and *Mertk* ^-/-V1^ mice were stained with hematoxylin and eosin. Representative retinal sections and quantification of ONL thickness in central (i), intermediate (ii) and peripheral (iii) retinal areas are shown. Scale bars = 200 μm (whole section) and 10 μm (insets). Data are represented as mean ± SEM of 15 measurements per area/ mouse, n = 3 mice/ genotype. Not significant (n.s.), *p<0.05, **p<0.01, ****p<0.0001 *vs*. C57BL/6 WT by 2-way ANOVA. **(C)** Eye transversal sections from P25, 6-month-old and 12-month old C57BL/6 WT and *Mertk* ^-/-V2^ mice were stained with hematoxylin and eosin. Representative retinal sections and quantification of ONL thickness in central (i), intermediate (ii) and peripheral (iii) retinal areas are shown. Scale bars = 200 μm (whole section) and 10 μm (insets). Data are represented as mean ± SEM of 15 measurements/ mouse, n = 3 mice/ genotype. Not significant (n.s.), *p < 0.05, **p < 0.01 *vs*. C57BL/6 WT by 2-way ANOVA. **(D)** Eye transversal sections from 12-month-old C57BL/6 WT and *Tyro3* ^-/-V1^ mice were stained with hematoxylin and eosin. Representative retinal sections and quantification of ONL thickness in central (i), intermediate (ii) and peripheral (iii) retinal areas are shown. Scale bars = 200 μm (whole section) and 10 μm (insets). Data are represented as mean ± SEM of 15 measurements/ mouse, n = 3 mice/genotype. Not significant (n.s.) *vs*. C57BL/6 WT by 2-way ANOVA.

We have recently generated an independent mouse line bearing *Mertk* ablation (termed *Mertk* ^-/-V2^ mice) ^22^. Histological analyses of the retina from *Mertk* ^-/-V2^ mice revealed that there were no statistically significant differences of the central and intermediate portions of the retina of *Mertk* ^-/-V2^ mice by comparison to B6 WT, at P25, 6 months or even at 12 months of age (**Fig. 1C**). Nonetheless, we did detect statistically significant differences between *Mertk* ^-/-V2^ and B6 WT mice in the peripheral retina (**Fig. 1C**). There was significant thinning of the peripheral retina, by comparison with B6 WT, at 6 and 12 months of age (**Fig. 1C**). Still, *Mertk* ^-/-V2^ mice did not display deficits in ERG responses even at 12 months of age (**Fig. S1A-D)**. Retinal histology and ERG responses were also preserved in *Tyro3* ^-/-V1^ mice even at 1 year of age (**Fig. 1D; Fig. S1A-D**).

Taken together, our findings are consistent with those of Vollrath *et al*. that demonstrated that in *Mertk* ^-/-^ *Tyro3* ^B6/129^ RPE, wherein TYRO3 expression was at 67% of *Tyro3* ^B6/B6^, the level of TYRO3 expression was enough to prevent severe, early-onset retinal degeneration ^17^. Severe, early-onset retinal degeneration was detected in *Mertk* ^-/-^ *Tyro3* ^B6/-^ (TYRO3 amounts at 50% of *Tyro3* ^B6/B6^) and progressively worsen in *Mertk* ^-/-^ *Tyro3* ^129/129^ (TYRO3 amounts at ∼33% of *Tyro3* ^B6/B6^) and in *Mertk* ^-/-^ *Tyro3* ^-/-17^. *Mertk* ^-/-^ *Tyro3* ^B6/129^ RPE did exhibit degeneration in the periphery ^17^, similar to our *Mertk* ^-/-V2^ mice. The age-related, progressive PR degeneration in *Mertk* ^-/-V2^ mice is also consistent with the observations of Maddox *et al*. wherein an ENU-mutagenesis that generated a homozygous *Mertk* ^nmf12^ mutation substituting a highly conserved histidine residue to arginine at amino acid position 716 within the tyrosine kinase domain and a 2.5-fold reduction in the amount of MERTK by immunoblotting, failed to phenocopy the severe, early-onset PR degeneration of *Mertk* ^-/-V1^ mice ^23^. Instead, *Mertk* ^*nmf12*^ mutated mice exhibited thinning of the peripheral ONL at P30, while the central retina was spared ^23^. ERG waveforms were detectable in *Mertk* ^*nmf12*^ homozygotes mice up to the age of 2 years ^23^. Thus, the loss of *Mertk* and the concomitant hypomorphic expression/simultaneous loss of *Tyro3* as in *Mertk* ^-/-V1^ and *Mertk* ^-/-V2^ *Tyro3* ^-/-V2^ mice, respectively, result in severe, early-onset PR degeneration. The loss of *Mertk* alone, as in *Mertk* ^-/-V2^ mice or the previously reported *Mertk* ^-/-^ *Tyro3* ^B6/129 17^ and *Mertk* ^*nmf12*^ mice ^23^ is associated with peripheral degeneration.

### Inflammation is detected in the retina of *Mertk* ^-/-V1^ and *Mertk* ^-/-V2^ *Tyro3* ^-/-V2^ mice

Audo *et al*. reported hyper-reflective dots above the RPE/Bruch’s membrane complex in all of their *MERTK* mutation-associated RP patients ^3^. These hyper-reflective dots were conjectured to be macrophages or activated microglia ^3^. Similarly, increased number of fluorescent spots indicative of activated microglia was reported in *Mertk* ^-/-V1^ mice, and IBA1^+^ cells were detected within the RPE and PR layers ^24^. Moreover, in the RCS rat, IBA1^+^ cells were detected in the ONL at P30 ^25^. By P60, an increased number of IBA1^+^ cells was detected across the entire retina, which decreased by P90 concurrently with the apoptosis of retinal cells ^25^. We have also previously reported that IBA1^+^ microglial numbers are ∼1.8 fold higher in the RCS rat than in WT rats between P14 to P30, and that RCS rat microglia migrate to the PR layer starting at P20 ^26^. Consistent with these observations, we noted the activation of resident immune cells and/or the infiltration of monocytes by flow cytometry analysis on the neural retina of *Mertk* ^-/-V1^, *Mertk* ^-/-V2^ *Tyro3* ^-/-V2^ and B6 WT mice. CD45^+(hi/int)^ cells were defined as CD11b^+^CD64^+^Ly6C^-^Ly6G^-^ microglia or CD11b^+^CD64^+^Ly6C^+^Ly6G^-^ monocytes. At P42, there was a significantly larger number of CD45^+^ cells in *Mertk* ^-/-V1^ and *Mertk* ^-/-V2^ *Tyro3* ^-/-V2^ retinas, relative to B6 WT retinas (**Fig. S2A, C**). We observed a statistically significant ∼2.5-fold increase of CD45^+^ cells in *Mertk* ^-/-V1^ retinas and ∼3-fold increase of CD45^+^ cells in *Mertk* ^-/-V2^ *Tyro3* ^-/-V2^ retinas, relative to B6 WT retinas. The number of CD45^int^CD11b^+^CD64^+^Ly6C^-^Ly6G^-^ microglia expanded significantly by ∼ 2.5-fold in *Mertk* ^-/-V1^ and 3-fold in *Mertk* ^-/-V2^ *Tyro3* ^-/-V2^ retinas, relative to B6 WT retinas (**Fig. S2A, C**). Similarly, CD45^hi^CD11b^+^CD64^-^Ly6C^+^Ly6G^-^ monocytes expanded significantly by ∼ 4-fold in *Mertk* ^-/-V1^ and 2-fold in *Mertk* ^-/-V2^ *Tyro3* ^-/-V2^ retinas, relative to B6 WT retinas (**Fig. S2A, C**). The microglia in *Mertk* ^-/-V1^ and *Mertk* ^-/-V2^ *Tyro3* ^-/-V2^ mice were activated, *i*.*e*. these cells expressed significantly higher amounts of activation markers such as CD68, CD11c and GAL3, when compared to cells from B6 WT mice (**Fig. S2B, D**).

To test whether microglial activation and/or monocyte infiltration was observed in *Mertk* ^-/-V1^ and *Mertk* ^-/-V2^ *Tyro3* ^-/-V2^ retinas even at a time-point in which retinal degeneration was histologically not yet observable, we performed similar flow cytometry analysis in the retina of P25 mice. Again, we detected a significant increase in the number of CD45 ^Int/+^ cells in the retinas of *Mertk* ^-/-V1^ (∼ 4-fold) and *Mertk* ^-/-V2^ *Tyro3* ^-/-V2^ (∼ 5-fold) *versus* B6 WT mice (**Fig. 2A, C**). The number of microglia was correspondingly increased by ∼3 fold in *Mertk* ^-/-V1^ and ∼ 8-fold in *Mertk* ^-/-V2^ *Tyro3* ^-/-V2^ *versus* B6 WT retinas (**Fig. 2A, C**). Moreover, the expression level of activation markers (CD68, CD11c and CD86) was also significantly elevated in these cells isolated from *Mertk* ^-/-V1^ and *Mertk* ^-/-V2^ *Tyro3* ^-/-V2^ retinas relative to B6 WT controls (**Fig. 2B, D**). Unlike at P42, CD45^hi^ cells were not detectable at P25 (**Fig. 2A, C**). In agreement with our flow cytometry findings at P25, immunofluorescence analysis of the total number of IBA1^+^ cells within the outer retina showed a significant increase in the number of IBA1^+^ cells in samples collected from P25 *Mertk* ^-/-V1^ and *Mertk* ^-/-V2^ *Tyro3* ^-/-V2^ mice relative to B6 WT controls (**Fig. 2E**). Furthermore, we observed increased translocation of microglia to the ONL layer in *Mertk* ^-/-V1^ and *Mertk*^-/-V2^ *Tyro3* ^-/-V2^ mice. By contrast, 100% of IBA1^+^ cells localized to the outer plexiform layer (OPL) in samples obtained from age-matched B6 WT controls (**Fig. 2E**). In addition to flow cytometry and immunohistological analysis, we employed Luminex^®^ multiplexed cytokine array analyses on P25 lysates of retina and RPE obtained from *Mertk* ^-/-V1^, *Mertk* ^-/-V2^ *Tyro3* ^-/-V2^ and B6 WT mice. This assay demonstrated elevated levels of the chemokine/cytokines CXCL10, CCL12, CCL19, MCSF and LIF in *Mertk* ^-/-V1^ and *Mertk* ^-/-V2^ *Tyro3* ^-/-V2^ lysates by comparison to their B6 WT counterparts (**Fig. 2F**). Collectively, these results indicate that the absence of *Mertk* together with the hypomorphic expression/loss of *Tyro3* is associated with increased production of inflammatory cytokines and chemokines, microglial activation, proliferation and translocation from the OPL to the ONL by P25, an age at which no histological signs of retinal degeneration were detected. Thus, inflammation is a feature of two independent mouse models of severe, early-onset PR degeneration. Furthermore, like in the RCS rat, inflammation is not simply a consequence of retinal degeneration but is detected before histological changes are observed in *Mertk* ^-/-V1^ and *Mertk* ^-/-V2^ *Tyro3* ^-/-V2^ retina.

**Fig. 2.**
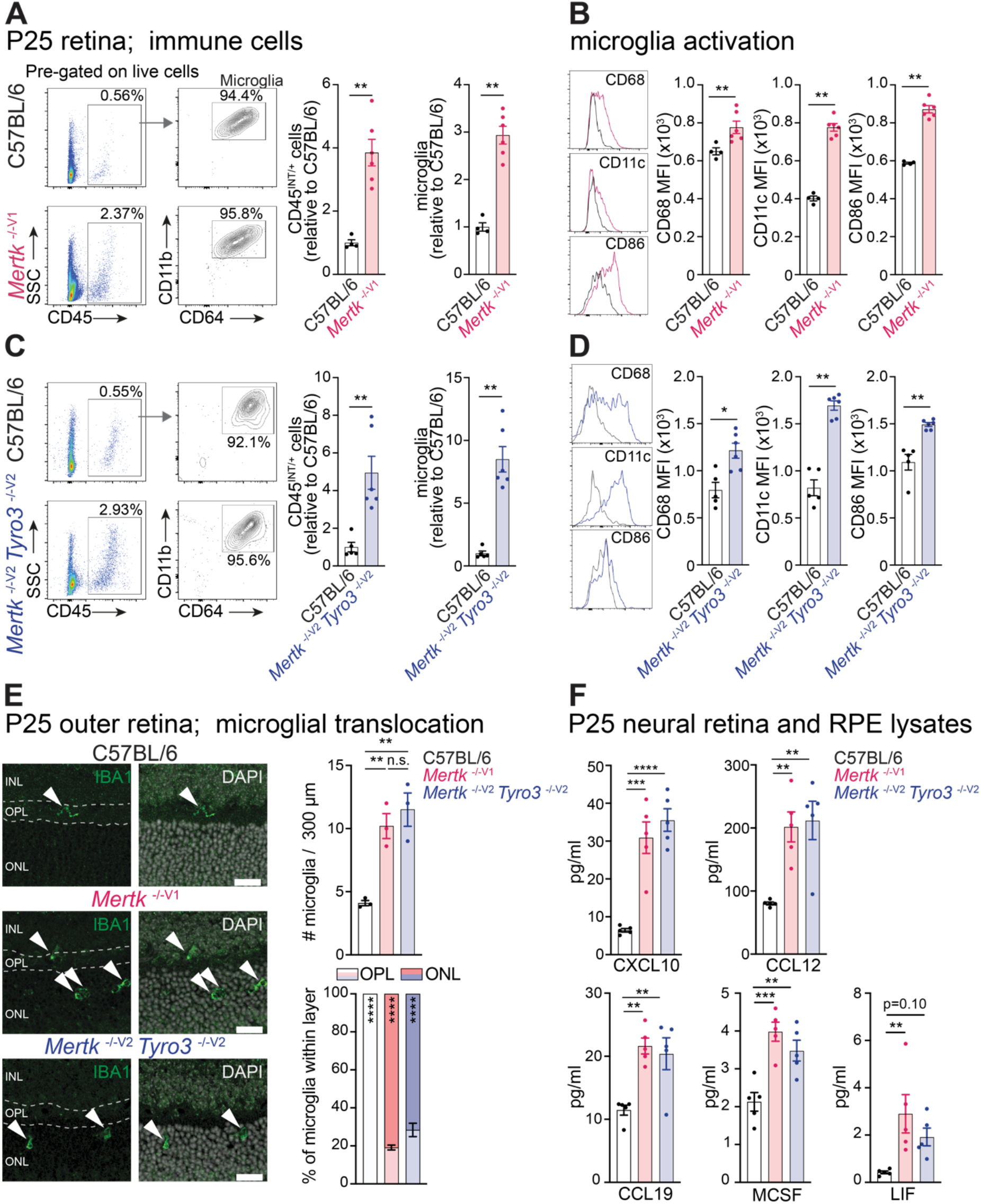
Loss of *Mertk* and hypomorphic expression or concomitant loss of *Tyro3* leads to retinal inflammation. **(A)** Retinas from P25 C57BL/6 WT and *Mertk* ^-/-V1^ were dissociated, and the type, number and activation of immune cells were determined by flow cytometry. The number of CD45^INT/+^ cells and microglia (CD45^INT/+^ CD11b^+^ CD64^+^ cells) within retinas are expressed as fold-change relative to the mean of P25 C57BL/6 WT and represented as mean ± SEM. N = 4-6 mice/ group. **p < 0.01 by Mann-Whitney’s test. **(B)** Representative histograms showing expression of CD68, CD11c and CD86 on microglia are shown, next to corresponding quantification of mean fluorescence intensity (MFI) represented as mean ± SEM. N = 4-6 mice/ group. **p < 0.01 by Mann-Whitney’s test. **(C)** Retinas from P25 C57BL/6 WT and *Mertk* ^-/-V2^ *Tyro3*^-/-V2^ were dissociated, and the type, number and activation of immune cells were determined by flow cytometry. The number of CD45^INT/+^ cells and microglia (CD45^INT/+^ CD11b^+^ CD64^+^ cells) within retinas are expressed as fold-change relative to the mean of P25 C57BL/6 WT and are represented as mean ± SEM. N = 5-6 mice/ group. **p < 0.01 by Mann-Whitney’s test. **(D)** Representative histograms showing expression of CD68, CD11c and CD86 on microglia are shown, next to corresponding quantification of mean fluorescence intensity (MFI). N = 5-6 mice/ group. *p<0.05, **p < 0.01 by Mann-Whitney’s test. **(E)** Representative IBA1 immunofluorescence staining, indicating microglia localization in retinas from P25 C57BL/6 WT, *Mertk* ^-/-V1^ and *Mertk* ^-/-V2^ *Tyro3*^-/-V2^ mice. Arrowheads indicate IBA1^+^ cells. Scale bars = 20 μm. Quantification of the total number of IBA1^+^ cells present within the outer retina (mean ± SEM, n = 3 mice/ group, not significant (n.s.), **p < 0.01 by one-way ANOVA-Tukey’s test). Quantification of the percentage of IBA1^+^ cells present in ONL *vs*. OPL (mean ± SEM, n = 3 mice/ group. ****p < 0.0001 *vs*. ONL, two-way ANOVA). INL: inner plexiform layer; OPL: outer plexiform layer; ONL: outer nuclear layer. **(F)** Quantification of levels of various chemokines and cytokines in lysates of P25 neural retinas and RPE of C57BL/6 WT, *Mertk* ^-/-V1^ and *Mertk* ^-/-V2^ *Tyro3* ^-/-V2^ mice by Luminex (mean ± SEM, n = 5 mice/ genotype). **p < 0.01, ***p < 0.001, ****p < 0.0001 *vs*. C57BL/6 WT by one-way ANOVA-Dunnet’s test.

### RPE inflammation is a distinct hallmark of *Mertk* ^-/-V1^ and *Mertk* ^-/-V2^ *Tyro3* ^-/-V2^ mice, but not a general feature of mice with genetic ablation of phagocytosis receptors

Albeit microglia are activated in the two models of severe, early-onset PR degeneration, both these models require the loss of *Mertk* together with the hypomorphic expression/loss of *Tyro3*. Microglia, however, express only *Mertk* and do not express *Tyro3*. Unlike microglia, RPE co-expresses *Mertk* and *Tyro3* ^16,17,27^. Therefore, we tested if there was an RPE-intrinsic inflammatory response in the *Mertk* ^-/-V1^ and *Mertk* ^-/-V2^ *Tyro3* ^-/-V2^ mice. We performed unbiased transcriptomics to compare the gene expression landscape of isolated RPE cells from B6 WT, *Mertk* ^-/-V1^ and *Mertk* ^-/-V2^ *Tyro3* ^-/-V2^ mice at P25 (**Fig. 3**). Pairwise comparison of RNA sequencing (RNAseq) data revealed 67 differentially expressed genes (DEGs) between *Mertk* ^-/-V1^ and B6 WT RPE (**Fig. 3A**). Interestingly, the differentially upregulated genes in *Mertk* ^-/-V1^ compared to B6 WT RPE corresponded to pathways related to innate immune response, response to IFN-α and IFN-γ, response to TNF-α, among other immune-related categories (**Fig. 3B**). To gain a broader understanding of the transcriptional changes in *Mertk* ^-/-V1^ *versus* B6 WT RPE, we performed gene set enrichment analysis (GSEA). This analysis also revealed pathways involved in the response to the inflammatory cytokines IFN-α, IFN-γ, IL-6 and TNF-α as top enriched categories in *Mertk* ^-/-V1^ RPE in comparison to B6 WT RPE (**Fig. 3C**). Similar to *Mertk* ^-/-V1^ RPE cells, we found 102 DEGs in *Mertk* ^-/-V2^ *Tyro3* ^-/-V2^ RPE when compared to B6 WT control RPE (**Fig. 3D**). The top enriched categories corresponding to the significantly upregulated genes in *Mertk* ^-/-V2^ *Tyro3* ^-/-V2^ RPE *versus* B6 WT RPE were immune response-related categories, similar to what was observed for the genes significantly upregulated in *Mertk* ^-/-V1^ RPE (**Fig. 3E**). Consistently, GSEA of RPE RNA-seq data identified genes associated with the response to IFN-α, IFN-γ, IL-6 and TNF-α to be significantly enriched in *Mertk* ^-/-V2^ *Tyro3* ^-/-V2^ RPE compared to B6 WT RPE (**Fig. 3F**). We validated the differential expression of inflammatory response signaling genes, such as *Cxcl10, Ifit1, Gm4951* and *Rsad2*, in *Mertk*^-/-V1^ and *Mertk* ^-/-V2^ *Tyro3* ^-/-V2^ RPE by qPCR (**Fig. 3G, H**). To confirm that these changes were indeed stemming primarily from RPE cells, and not from contaminating immune cells, we determined the expression of candidate RPE-specific *versus* immune cell-specific genes in our dataset. We found that compared to counts for pan-immune cell marker CD45 (*Ptprc*), macrophage markers F4/80 (*Adgre1*) and CD11b (*Itgam*), and microglia markers CX3CR1 (*Cx3rcr1*) and IBA-1 (*Aif1*), RPE genes *Rpe65, Ezr, Rlbp1* were enriched by >200-fold (**Fig. 3I**). This enrichment of RPE genes allowed us to attribute the inflammatory module observed in *Mertk* ^-/-V1^ and *Mertk* ^-/-V2^ *Tyro3* ^-/-V2^ mice, primarily to RPE cells. In addition to RNA sequencing analysis, we employed Luminex^®^ multiplexed cytokine array analyses on P25 lysates of RPE obtained from *Mertk* ^-/-V1^, *Mertk* ^-/-V2^ *Tyro3* ^-/-V2^ and B6 WT mice. This assay demonstrated elevated levels of the chemokine/cytokines CXCL10, CCL12, CCL19 and IL-5 in *Mertk* ^-/-V1^ and *Mertk* ^-/-V2^ *Tyro3* ^-/-V2^ RPE lysates by comparison to their B6 WT counterparts (**Fig. 3J**). Thus, an RPE-intrinsic inflammation is a feature of the severe, early-onset PR degeneration.

**Fig. 3.**
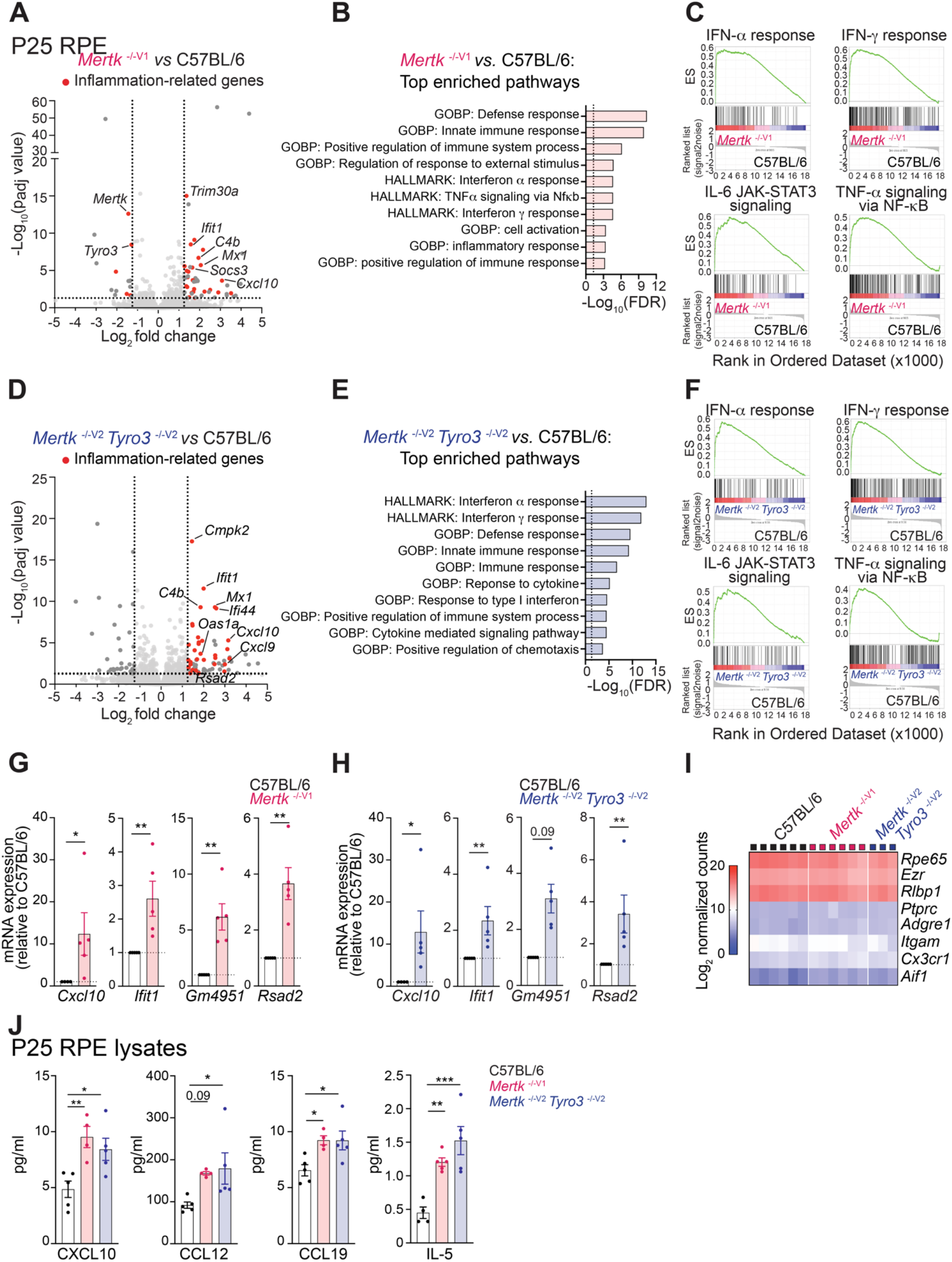
Loss of *Mertk* and hypomorphic expression or concomitant loss of *Tyro3* triggers exacerbated inflammation in the RPE. **(A)** Volcano plot comparing gene expression at P25 in *Mertk* ^-/-V1^ RPE relative to C57BL/6 WT RPE by RNA sequencing (n = 6 samples/ genotype, each sample was comprised of pooled tissues from 2 animals). Dashed lines indicate thresholds for significance (FC>2 or FC<-2 and padj<0.05). Genes indicated in red have previously been associated with inflammation. **(B)** Top enriched Gene Ontology – Biological Process (GOBP) and hallmark pathways comprised of genes significantly upregulated at P25 in *Mertk* ^-/-V1^ RPE relative to C57BL/6 WT RPE. Dashed line indicates threshold for significance (FDR<0.05) **(C)** Gene set enrichment analysis comparing *Mertk* ^-/-V1^ *vs*. C57BL/6 WT -derived RPE. Significantly upregulated categories related to inflammation in *Mertk* ^-/-V1^ mice are shown. ES, enrichment score. **(D)** Volcano plot comparing gene expression at P25 in *Mertk* ^-/-V2^ *Tyro3* ^-/-V2^ RPE relative to C57BL/6 WT RPE by RNA sequencing (n = 3-6 samples/ genotype, each sample was comprised of pooled tissues from 2 animals). Dashed lines indicate thresholds for significance (FC>2 or FC<-2 and padj<0.05). Genes indicated in red have previously been associated with inflammation. **(E)** Top enriched GOBP and hallmark pathways comprised of genes significantly upregulated at P25 in *Mertk* ^-/-V2^ *Tyro3* ^-/-V2^ RPE relative to C57BL/6 WT RPE. Dashed line indicates threshold for significance (FDR<0.05) **(F)** Gene set enrichment analysis comparing *Mertk* ^-/-V2^ *Tyro3* ^-/-V2^ *vs*. C57BL/6 WT-derived RPE. Significantly upregulated categories related to inflammation in *Mertk* ^-/-V2^ *Tyro3*^-/-V2^ mice are shown. ES, enrichment score. **(G) (H)** qPCR validation of inflammation-associated genes upregulated in *Mertk* ^-/-V1^ (**G**) or *Mertk* ^-/-V2^ *Tyro3*^-/-V2^ (**H**) *vs*. C57BL/6 WT RPE (mean ± SEM, n = 5 mice/ genotype). *p < 0.05, **p < 0.01 by Mann-Whitney’s test. **(I)** Heatmap indicating log_2_ normalized counts of RPE and immune cell-associated genes at P25 in C57BL/6 WT, *Mertk* ^-/-V1^ and *Mertk* ^-/-V2^ *Tyro3* ^-/-V2^ RPE by RNA sequencing. **(J)** Quantification of levels of various chemokines and cytokines in lysates of P25 RPE of C57BL/6 WT, *Mertk* ^-/-V1^ and *Mertk* ^-/-V2^ *Tyro3* ^-/-V2^ mice by Luminex (mean ± SEM, n = 4-5 mice/ genotype). *p<0.05, **p < 0.01, ***p < 0.001 *vs*. C57BL/6 WT by one-way ANOVA-Dunnet’s test.

What leads to RPE inflammation in *Mertk* ^-/-V1^ and *Mertk* ^-/-V2^ *Tyro3* ^-/-V2^ mice? PR degeneration in *Mertk* ^-/-V1^ mice has historically been associated with distortion of PR outer segments and subretinal debris buildup due to a drastic reduction in POS uptake. Thus, *a priori*, RPE inflammation may be thought of as an obligate consequence of reduced phagocytosis. Instead, we posit that RPE inflammation distinct from deficient phagocytosis drives severe, early-onset PR degeneration. Given that *Mertk* ^-/-V2^ mice do not phenocopy the severe, early-onset inflammation, we hypothesized that it would not display RPE inflammation despite defective phagocytosis. To quantitate phagocytosis, we counted the number of rhodopsin-positive phagosomes in randomly chosen fields across all regions within RPE flat mounts in *Mertk* ^-/-V2^ and B6 WT mice. Phagosomes were quantified 1 hour after light onset, when phagosome load in B6 WT RPE is at its peak. The diameter of a POS is estimated to be ∼1 μm ^28^. Therefore, we quantified the number of rhodopsin-positive phagosomes with diameter ≥ 0.9 μm within RPE cells as a proxy for phagosomes that contain undigested, and thus most likely recently engulfed, outer segment tips. Additionally, we counted rhodopsin-positive phagosomes ≥ 0.5 μm in diameter for phagosomes containing intact as well as digested or partially digested outer segment tips. We determined the level of phagocytosis at P25, an age when retinal histology was yet unaffected (**Fig. 1**). The mean ± SEM from B6 WT RPE was 41.9 ± 4.3 undigested ≥ 0.9 μm POS phagosomes (**Fig. 4A**). *Mertk* ^-/-V2^ RPE, by contrast, had 24.6 ± 3.0 undigested POS phagosomes (**Fig. 4A**). Quantification of all rhodopsin-positive phagosomes ≥ 0.5 μm in diameter revealed that while B6 WT RPE had 168.1 ± 4.31 rhodopsin-positive phagosomes, in *Mertk* ^-/-V2^ RPE the number of phagosomes were 113.1 ± 7.63. When considering ≥ 0.5 μm phagosomes, we noted that their numbers within the *Mertk* ^-/-V2^ RPE were heterogenous within an individual eye, with the number of phagosomes ranging from a maximum of 261.68/ field to a lowest of 34.09/field. By contrast, a maximum of 259.20/ field to a minimum of 96.87/ field were observed in B6 WT RPE (**Fig. 4A**).

**Fig. 4.**
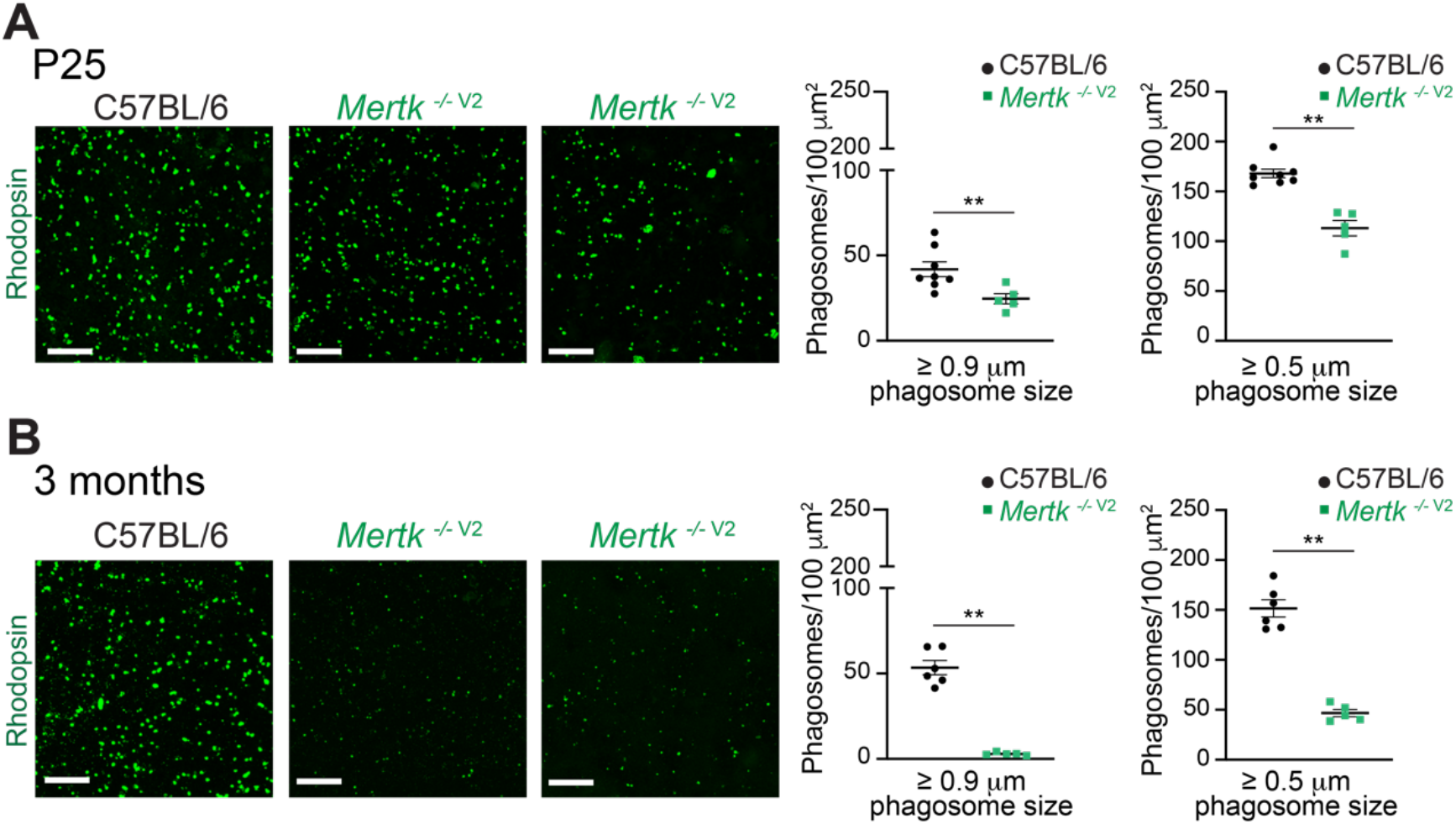
Loss of *Mertk* results in early-onset defects in RPE POS phagocytosis *in vivo*. **(A)** POS phagocytosis was analyzed by counting rhodopsin-positive inclusions in whole-mount RPE from P25 C57BL/6 WT and *Mertk* ^-/-V2^ mice sacrificed 1 hour after light onset. Representative images, large phagosome counts (recently engulfed POS, diameter ≥0.9μm) and total phagosome counts (diameter ≥0.5μm) are shown. Data are mean ± SEM of 13-16 fields/ mouse, n = 5 - 8 mice/ genotype. **p < 0.01 *vs*. C57BL/6 WT by Mann-Whitney’s test. Scale bars = 20 μm. **(B)** POS phagocytosis was analyzed by counting rhodopsin-positive inclusions in whole-mount RPE from 3-month-old C57BL/6 WT and *Mertk* ^-/-V2^ mice sacrificed 1 hour after light onset. Representative images, large phagosome counts (recently engulfed POS, diameter ≥0.9μm) and total phagosome counts (diameter ≥0.5μm) are shown. Data are mean ± SEM of 16 fields/ mouse, n = 5 - 6 mice/ genotype. **p < 0.01 *vs*. C57BL/6 WT by Mann-Whitney’s test. Scale bars = 20 μm.

We also tested the *Mertk* ^-/-V2^ mice for engulfment of rhodopsin-positive phagosomes at 3 months of age. B6 WT RPE contained 53.4 ± 4.19 undigested ≥ 0.9 μm POS phagosomes at 1 hour after light onset (**Fig. 4B**). In stark contrast, *Mertk* ^-/-V2^ RPE had only 2.93 ± 0.395 undigested POS phagosomes. Similarly, rhodopsin-positive phagosomes ≥ 0.5 μm in diameter were 152 ± 8.63 in B6 WT RPE, but only 46.8 ± 3.66 in *Mertk* ^-/-V2^ RPE (**Fig. 4B**). At 3 months of age, the numbers of phagosomes/field in *Mertk* ^-/-V2^ RPE were no longer heterogenous, consistent with a progressive phagocytosis deficiency worsening from P25 to 3 months of age. The size and distribution of *Mertk* ^-/-V2^ RPE cells were indistinguishable from B6 WT controls at both P25 and 3 months of age as assessed by F-actin staining (**Fig. S3**). Collectively, our results demonstrate a significant reduction in phagocytosis of outer segment tips by the RPE in *Mertk* ^-/-V2^ mice in comparison to RPE in B6 WT mice.

Despite this deficit in phagocytosis, *Mertk* ^-/-V2^ retina did not exhibit expansion of CD45^+(hi/int)^ cells, nor proliferation or activation of CD45^int^CD11b^+^CD64^+^ microglia or infiltration of CD45^hi^ monocytes at P25 (**Fig. 5A**). The number of CD45^+(hi/int)^ cells were statistically no different in *Mertk* ^-/-V2^ retina than in B6 WT retina, and unlike the expansion and activation observed in *Mertk* ^-/-V2^ *Tyro3* ^-/-V2^ retina (**Fig. 5A**). Correspondingly, there was no statistically significant change in microglial activation marker expression in *Mertk* ^-/-V2^ retina relative to B6 WT retina (**Fig. 5B**). Luminex^®^ multiplexed cytokine array analyses did not show enhanced amounts of CXCL10, CCL12, CCL19, MCSF and LIF in lysates from *Mertk* ^-/-V2^ neural retina and RPE as detected in *Mertk* ^-/-V1^ and *Mertk* ^-/-V2^ *Tyro3* ^-/-V2^ retina and RPE at P25 (**Fig. 5C**). Only CCL19 was higher in the *Mertk* ^-/-V2^ retina-RPE lysate than that in B6 WT, albeit not to the same level as in *Mertk* ^-/-V1^ and *Mertk* ^-/-V2^ *Tyro3* ^-/-V2^ retina and RPE, while all other cytokines/chemokines in *Mertk* ^-/-V2^ retina-RPE lysate were not significantly different that in B6 WT neural retina and RPE (**Fig. 5C**). GSEA of RPE RNA-seq data and qPCR analyses revealed that *Mertk* ^-/-V2^ mice failed to exhibit increased inflammation as *Mertk* ^-/-V1^ and *Mertk* ^-/-V2^ *Tyro3* ^-/-V2^ RPE at P25 (**Fig. 5D**). The *Mertk* paralog *Tyro3* was shown to have a role in RPE phagocytosis *in vitro* ^17^. qRT-PCR validation of *Ifit1, Iigp1* and *Rsad2* confirmed the lack of heightened inflammatory response in *Mertk* ^-/-V2^ or *Tyro3* ^-/-V1^ RPE relative to C57BL/6, and distinctly opposed to the exacerbated inflammatory response in *Mertk* ^-/-V1^ or *Mertk* ^-/-V2^ *Tyro3* ^-/-V2^ RPE (**Fig. 5F**). We also examined features of inflammation in *Tyro3* ^-/-V1^ retina and failed to detect any significant changes compared to B6 WT (**Fig. 5A-C, E, F**).

**Fig. 5.**
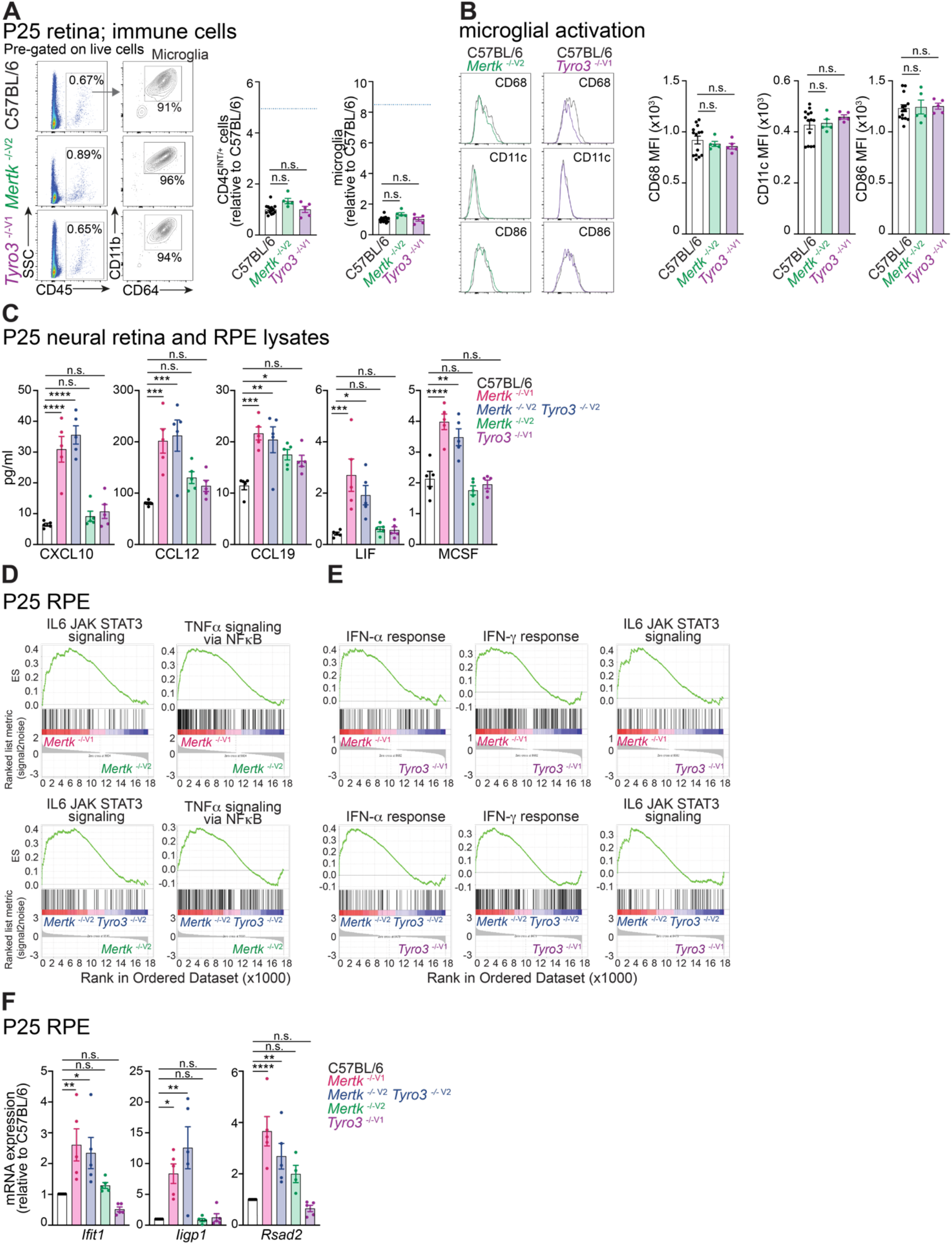
Independent ablation of *Mertk* or *Tyro3* does not lead to retinal inflammation. **(A)** Retinas from P25 C57BL/6 WT, *Mertk* ^-/-V2^ and *Tyro3* ^-/-V1^ mice were dissociated, and the number, type and activation of immune cells were determined by flow cytometry. Number of CD45^INT/+^ cells and microglia (CD45^INT/+^ CD11b^+^ CD64^+^ cells) are expressed as fold-change relative to the mean of P25 C57BL/6 WT (mean ± SEM, n = 5-14 mice/ group, not significant (n.s.) by one-way ANOVA). Dashed line indicates the fold change of CD45^INT/+^ cells and microglia observed in *Mertk* ^-/-V2^ *Tyro3* ^-/-V2^ *vs*. C57BL/6 WT retinas at P25 (Corresponding to data in Figure 2C). **(B)** Representative histograms showing expression of CD68, CD11c and CD86 on microglia at P25 in C57BL/6 WT, *Mertk* ^-/-V2^ and *Tyro3* ^-/-V1^ mice are shown, next to corresponding quantification of mean fluorescence intensity (MFI) represented as mean ± SEM, n = 5-14 mice/ group, not significant (n.s.) by one-way ANOVA. **(C)** Quantification of amounts of chemokines and cytokines in the lysates of P25 neural retinas and RPE at P25 in C57BL/6 WT, *Mertk* ^-/-V1^, *Mertk* ^-/-V2^ *Tyro3*^-/-V2^, *Mertk* ^-/-V2^ and *Tyro3* ^-/-V1^ mice by Luminex. Data are represented as mean ± SEM, n = 5 mice/ genotype, not significant (n.s.), *p < 0.05, **p < 0.01, ***p < 0.001. ****p < 0.0001 *vs*. C57BL/6 WT by one-way ANOVA, followed by Dunnet’s test. Datasets from *Mertk* ^-/-V1^ and *Mertk* ^-/-V2^ *Tyro3* ^-/-V2^ mice correspond to data in Figure 2F. All measurements were done in parallel. **(D)** Gene set enrichment analysis comparing *Mertk* ^-/-V1^ *vs. Mertk* ^-/-V2^-derived RPE (top) or *Mertk* ^-/-V2^ *Tyro3* ^-/-V2^ *vs. Mertk* ^-/-V2^-derived RPE (bottom) at P25. ES, enrichment score. **(E)** Gene set enrichment analysis comparing *Mertk* ^-/-V1^ *vs. Tyro3* ^-/-V1^-derived RPE (top) or *Mertk* ^-/-V2^ *Tyro3* ^-/-V2^ *vs. Tyro3* ^-/-V1^-derived RPE (bottom) at P25. ES, enrichment score. **(F)** qPCR validation of inflammation-associated genes in RPE of P25 *Mertk* ^-/-V1^, *Mertk* ^-/-V2^ *Tyro3*^-/-V2^, *Mertk* ^-/-V2^ and *Tyro3* ^-/-V1^ mice, relative to C57BL/6 WT mice. Data are represented as mean ± SEM, n = 5 mice/ genotype. Not significant (n.s.), *p < 0.05, **p < 0.01, ****p < 0.0001 by one-way ANOVA, followed by Dunnet’s test.

*Itgb5* ^-/-^ mice provide an alternative model for studying the consequences of POS uptake defects. We previously reported examination of *in vivo* phagocytosis in 3–4-months old mice which revealed the complete absence of the burst of POS phagocytosis after light onset in *Itgb5* ^-/-^ RPE that is characteristic to RPE in WT mice in response to the circadian shedding of POS ^14^. *In vitro* phagocytosis assays confirmed a defect in POS recognition and thus POS engulfment in primary cultures of RPE cells from *Itgb5* ^-/-^ mice ^14^. We therefore examined the whole-genome transcriptional profile of *Itgb5* ^-/-^ RPE by RNA-seq. We found that expression of the paired *Itgb5* and *Itgav* subunits were detected in RPE as early as P10 (**Fig S4A**). Unlike in *Mertk* ^-/-V2^ *Tyro3* ^-/-V2^ and *Mertk* ^-/-V1^ RPE, no inflammatory gene expression profile was observed in *Itgb5* ^-/-^ mice as compared to control WT^129^ mice at P25 (**Fig. S4B, C**). Taken together, these results indicate that phagocytic defects in RPE and reduced POS uptake do not obligatorily result in the RPE inflammation associated with severe, early-onset PR degeneration. Instead, RPE inflammation was exclusively associated with the genetic deletion of *Mertk* and the concomitant hypomorphic expression/simultaneous ablation of *Tyro3*, and not with the genetic deletion of *Mertk* alone or of *Tyro3* alone.

### Upregulation of inflammation-associated gene module in *Mertk* ^-/-V1^ and *Mertk* ^-/-V2^ *Tyro3* ^-/-V2^ RPE precedes eye opening

We next investigated if RPE inflammation can be temporally resolved from deficient phagocytosis in *Mertk* ^-/-V1^ and *Mertk* ^-/-V2^ *Tyro3* ^-/-V2^ mice. Phagocytosis of POS tips is minimal in rodents before eye-opening at P10 ^26^. We first examined whether *Mertk* and *Tyro3* expression is detected at P10, before eye-opening. We observed that both *Mertk* and *Tyro3* are expressed in B6 WT RPE, as detected by qPCR, at P10 (**Fig. 6A**). Next, we analyzed the expression of inflammation-associated genes at this developmental stage by qPCR. Remarkably, even at P10, *Mertk* ^-/-V1^ and *Mertk* ^-/-V2^ *Tyro3* ^-/-V2^ RPE cells were found to upregulate expression of genes involved in the response to pro-inflammatory cytokines IFN-α, IFN-γ, and TNF-α, by comparison with B6 WT RPE (**Fig. 6B, C**). Namely, we observed an ∼1.5 to 2.5-fold change in the expression of ISGs, including *Gm4951, Irf7, Irf9, Rsad2, Ifit3* and *Mx1* in *Mertk*^-/-V1^ and *Mertk* ^-/-V2^ *Tyro3* ^-/-V2^ RPE relative to B6 WT RPE (**Fig. 6B, C**). Similarly, TNF-α -response genes such as *Lif* and *Tnfsf9* were upregulated ∼1.5 to 2-fold in *Mertk* ^-/-V1^ and *Mertk* ^-/-V2^ *Tyro3* ^-/-V2^ RPE compared to B6 WT RPE (**Fig. 6B, C**). We also detected significantly increased expression (∼2 to 4.5-fold) of genes encoding for the chemokines *Ccl2 and Ccl3* in *Mertk* ^-/-V1^ and *Mertk* ^-/-V2^ *Tyro3* ^-/-V2^ RPE relative to B6 WT RPE (**Fig. 6B, C**). These results demonstrate that RPE inflammation precedes eye-opening. The early onset of RPE inflammation in the absence of *Mertk* and reduced expression/absence of *Tyro3* is thus unlikely to be a consequence of accumulated damage in the retina as a corollary of failed RPE phagocytosis.

**Fig. 6.**
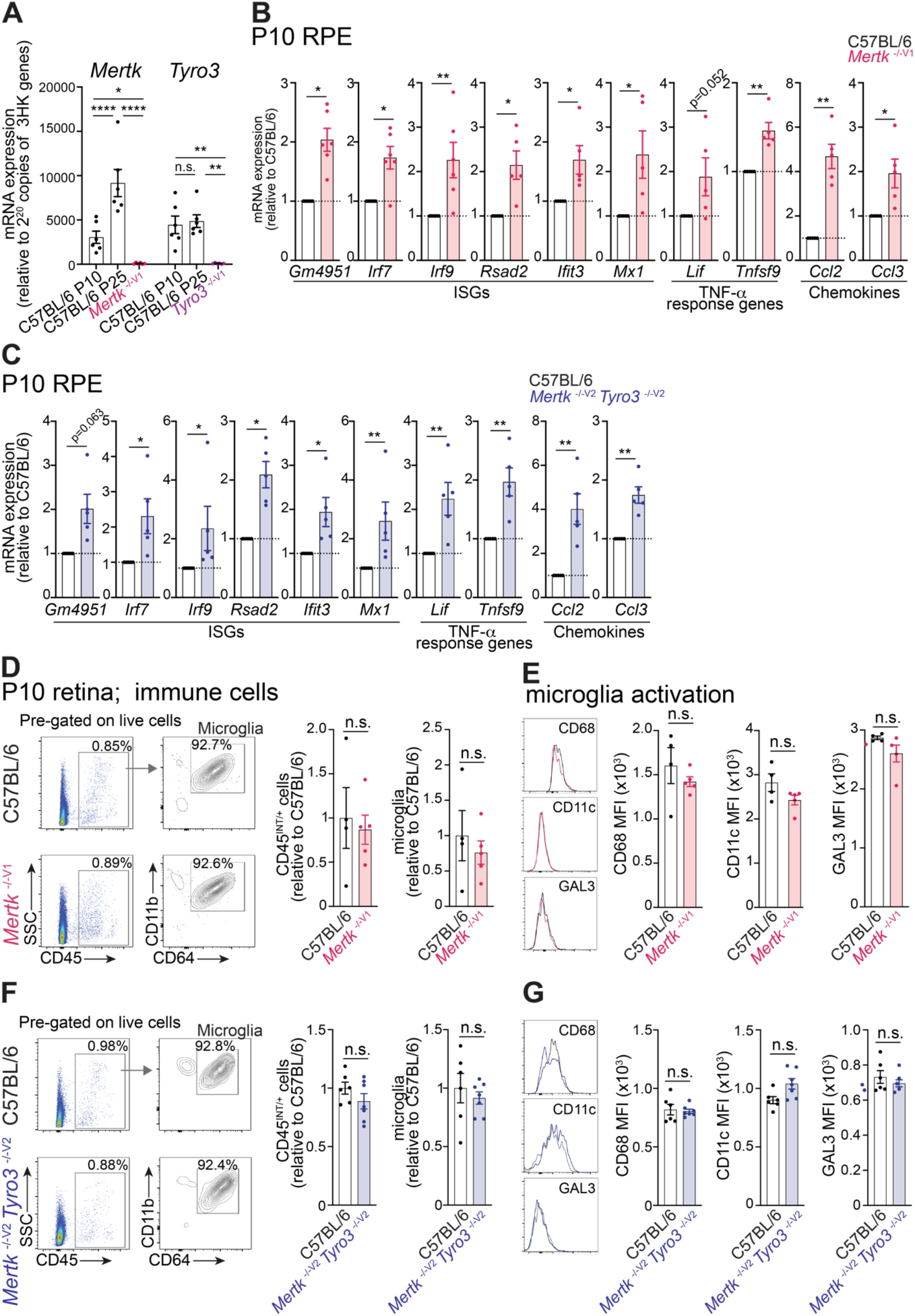
Increased inflammation in the RPE of *Mertk* ^-/-V1^ and *Mertk* ^-/-V2^ *Tyro3* ^-/-V2^ mice precedes eye opening and retinal microglia activation. **(A)** qPCR measurement of amounts of *Mertk* and *Tyro3* mRNA expression at P10 and P25 in C57BL/6 WT RPE (mean ± SEM, n = 6 mice/ group, not significant (n.s.), *p < 0.05, **p < 0.01, ****p < 0.0001 by one-way ANOVA, followed by Tukey’s test). **(B)** Assessment of interferon-stimulated genes (ISGs), TNF-α-inducible genes, and chemokines at P10 in RPE from C57BL/6 WT and *Mertk* ^-/-V1^ mice by qPCR (mean ± SEM, n = 5-6 mice/ group, *p < 0.05, **p < 0.01 by one-tailed Mann-Whitney’s test). **(C)** Assessment of ISGs, TNF-α-inducible genes, and chemokines at P10 in the RPE from C57BL/6 WT and *Mertk* ^-/-V2^ *Tyro3* ^-/-V2^ mice by qPCR (mean ± SEM, n = 5 mice/ group, *p < 0.05, **p < 0.01 by one-tailed Mann-Whitney’s test). **(D)** Retinas from P10 C57BL/6 WT and *Mertk* ^-/-V1^ mice were dissociated, and the type, number and activation of immune cells were determined by flow cytometry. Number of CD45^INT/+^ cells and of microglia (CD45^INT/+^ CD11b^+^ CD64^+^ cells) at P10 in C57BL/6 WT and *Mertk* ^-/-V1^ mice. Data are normalized to the mean of P10 C57BL/6 WT and represented as mean ± SEM, n = 4-5 mice/ group, not significant (n.s.), by Mann-Whitney’s test. **(E)** Representative histograms and corresponding mean fluorescence intensity (MFI) showing expression of CD68, CD11c and GAL3 on microglia in P10 *Mertk* ^-/-V1^ mice, as compared to age-matched C57BL/6 WT controls are shown, next to corresponding quantification of mean fluorescence intensity (MFI) represented as mean ± SEM. N =4 -5 mice/ group, not significant (n.s.) by Mann-Whitney’s test. **(F)** Retinas from P10 C57BL/6 WT and *Mertk* ^-/-V2^ *Tyro3* ^-/-V2^ mice were dissociated, and the type, number and activation of immune cells were determined by flow cytometry. Number of CD45^INT/+^ cells and of microglia (CD45^INT/+^ CD11b^+^ CD64^+^ cells) in C57BL/6 WT and *Mertk* ^-/-V2^ *Tyro3* ^-/-V2^ mice. Data are normalized to the mean C57BL/6 WT and represented as mean ± SEM, n = 6-7 mice/ group, not significant (n.s.), by Mann-Whitney’s test. **(G)** Representative histograms and corresponding mean fluorescence intensity (MFI) showing expression of CD68, CD11c and GAL3 on microglia in P10 *Mertk* ^-/-V2^ *Tyro3* ^-/-V2^ mice, as compared to age-matched C57BL/6 WT controls are shown, next to corresponding quantification of mean fluorescence intensity (MFI) represented as mean ± SEM. N = 6-7 mice/ group, not significant (n.s.) by Mann-Whitney’s test.

### RPE inflammatory response precedes microglial activation and subsequently cascades to involve other immune cells

The order in which an inflammatory response occurs in RPE, microglia and monocytes during severe, early-onset PR degeneration remains unclear. For example, microglial activation and consequent contribution to PR degeneration was shown in the RCS rat ^26^. However, it was also reported that the transcriptional reprogramming of retinal microglia from a homeostatic gene expression profile to the upregulation of injury response genes, in fact, confers protection to the RPE in models of acute light damage or the autosomal dominant knock-in rhodopsin mutation model of chronic damage ^29^. Therefore, microglial activation due to PR damage may be protective for RPE, but RPE inflammation causing microglial activation may in fact cause PR degeneration. In order to identify the nucleating event for retinal inflammation associated with severe, early-onset PR degeneration, we determined the temporal sequence of microglial *versus* RPE inflammation. We tested whether changes in microglial response in the neural retina were also already observed at P10 simultaneously with RPE inflammation, or if inflammatory changes were unique to RPE cells at this early age. We collected retinal samples from mice at P10 and performed flow cytometry analysis for microglia. Interestingly, we did not find a significant difference in the number of CD45 ^Int/+^ cells and of microglia in the neural retinas of *Mertk* ^-/-V1^ and *Mertk*^-/-V2^ *Tyro3* ^-/-V2^ mice, compared to B6 WT controls at P10 (**Fig. 6D, F**). Moreover, the expression level of markers associated with activation (CD68, CD11c and GAL3) was statistically no different in microglia isolated from *Mertk* ^-/-V1^ and *Mertk* ^-/-V2^ *Tyro3* ^-/-V2^ retinas relative to B6 WT controls (**Fig 6E, G**). Thus, at P10, the inflammatory response was confined to the RPE.

Taken together, our data suggests that inflammation is already present at P10, but it is restricted to the RPE. At P25, not only is inflammation detectable in the RPE, but it also extends beyond the RPE to manifest as expansion of microglial numbers and activation. By 6 weeks, unrestrained inflammation in the RPE and retinal microglia, and the resulting damage, leads to infiltration of monocytes into the retina. Thus, inflammation and retinal damage may constitute a feedback loop that perpetuate and exacerbate each other.

### Loss of *Mertk* function in microglia, with preserved *Mertk* and *Tyro3* in RPE, does not result in severe, early-onset PR degeneration

Our temporal assessment of the retinal inflammatory response could not unequivocally rule out the possibility that subthreshold changes in the microglia in the absence of *Mertk* may precipitate RPE inflammation at P10. Additionally, failed POS uptake due to the absence of microglial phagocytosis in *Mertk* ^-/-V1^ and *Mertk* ^-/-V2^ *Tyro3* ^-/-V2^ mice may drive RPE inflammation, as well as the inflammatory response of microglia and monocytes, in these mice. Although the loss of *Mertk* alone does not culminate in severe, early-onset PR degeneration as would be anticipated for a primarily microglia-driven phenotype, we attempted to definitively rule out the role of microglia in being the sole driver of the *Mertk*^-/-V1^ and *Mertk* ^-/-V2^ *Tyro3* ^-/-V2^ mice phenotype. First, we confirmed MERTK expression in microglia isolated from the retina by flow cytometry. Albeit at lower levels of expression than in brain microglia, retinal microglia indeed expressed an ample amount of MERTK (**Fig. 7A**). Next, we generated mice bearing conditional ablation of *Mertk* in myeloid, *Csf1r* ^+^ cells (*Csf1r-cre* ^*+*^ *Mertk* ^f/f^ mice). We confirmed that MERTK amounts were significantly reduced in *Csf1r-cre* ^*+*^ *Mertk* ^f/f^ mice when compared to littermate *Csf1r-cre* ^*-*^ *Mertk* ^f/f^ mice (**Fig. S5**). We then collected retinas from these mice at P25 and performed flow cytometry analysis to test whether the conditional genetic ablation of *Mertk* altered the number and/or the activation state of microglia. We found no significant changes in the number of CD45 ^Int/+^ cells and of microglia in the neural retinas of *Csf1r-cre* ^*+*^ *Mertk* ^f/f^ mice, compared to *Csf1r-cre* ^-^ *Mertk* ^f/f^ littermate controls (**Fig. 7B**). Additionally, the expression level of markers associated with activation (CD68, CD11c, and CD86) was statistically unchanged in *Csf1r-cre* ^*+*^ *Mertk* ^f/f^ microglia, compared to *Csf1r-cre* ^-^ *Mertk* ^f/f^ controls (**Fig. 7C**). Furthermore, histological analysis of eyes obtained from 12-months old *Csf1r-cre* ^*+*^ *Mertk* ^f/f^ mice showed preserved histoarchitecture of all retinal layers (**Fig. 7D**). Collectively, our results show that the loss of microglia-intrinsic function of *Mertk* with intact *Mertk* and *Tyro3* in the RPE does not give rise to early-onset retinal degeneration, in contrast to what was observed in *Mertk* ^-/-V2^ *Tyro3* ^-/-V2^ or *Mertk* ^-/-V1^ mice. Loss of *Mertk* in the microglia does not drive microglial activation in the absence of RPE inflammation. Therefore, the loss of MERTK function in microglia is either dispensable for the retinal degeneration phenotype, or at least is not sufficient for inducing retinal degeneration. A microglia-intrinsic function of this RTK remains a formal possibility, if it is to amplify a response initiating outside microglia, such as in the RPE.

**Fig. 7.**
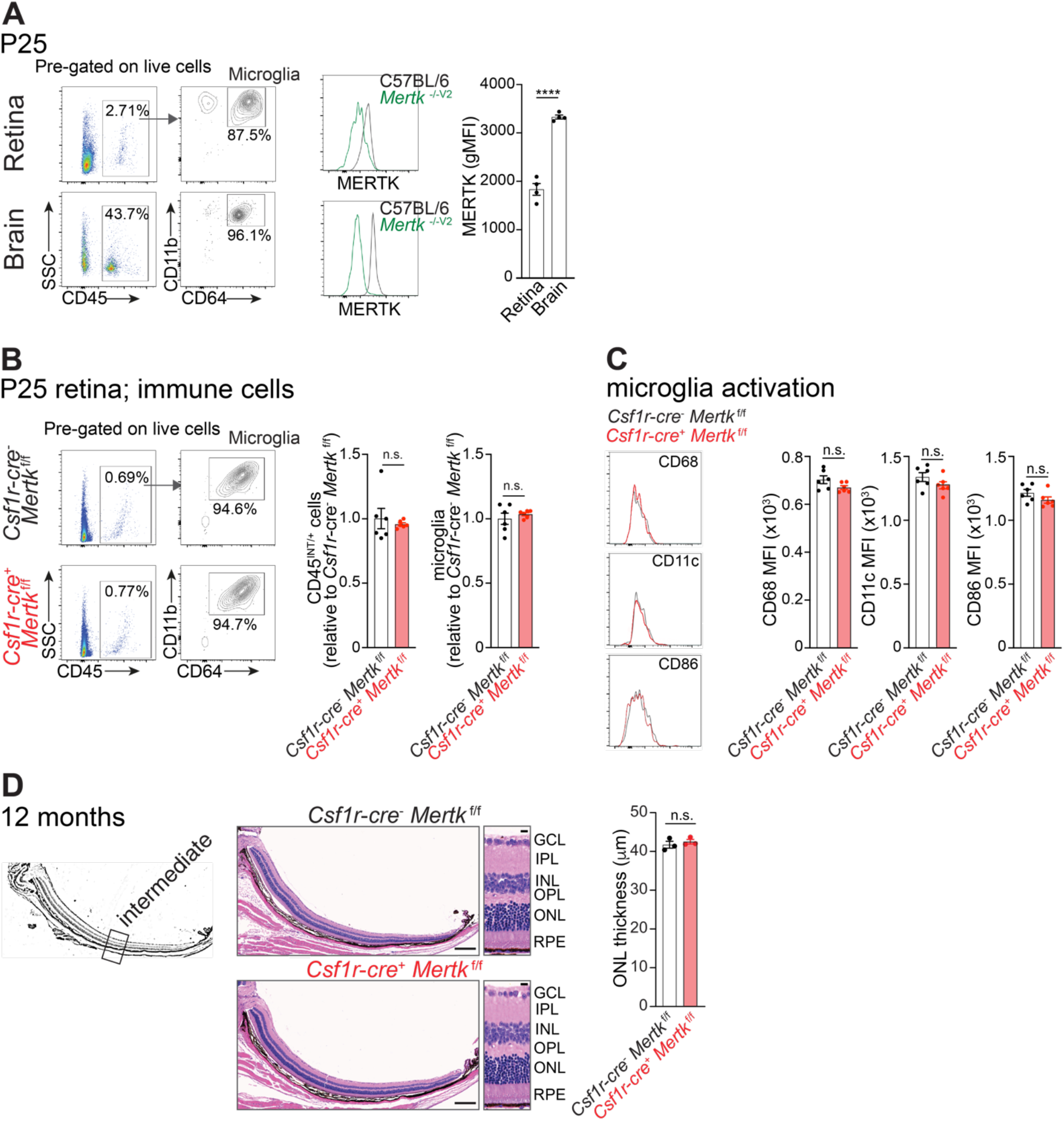
Conditional ablation of *Mertk* in microglia is not sufficient to drive retinal degeneration. **(A)** Retina and brain tissue were isolated from P25 C57BL/6 WT and *Mertk* ^-/-V2^ mice and stained for MERTK expression via flow cytometry. Gating strategy for identification of microglia (CD45^INT/+^ CD11b^+^ CD64^+^ cells) in the retina (upper panels) and the brain (lower panels), as well as representative histograms of MERTK expression on the surface of retina and brain microglia are shown, next to quantification of mean fluorescence intensity (MFI) of MERTK. Data are represented as mean ± SEM, n = 4 mice/ group, ****p < 0.0001 by Mann-Whitney’s test. **(B)** Retinal tissue was isolated at P25 from *Csf1r-cre*^*+*^ *Mertk* ^f/f^ and *Csf1r-cre*^*-*^ *Mertk* ^f/f^ mice. The number of CD45^INT/+^ cells and the number and activation of microglia (CD45^INT/+^ CD11b^+^ CD64^+^ cells) were determined by flow cytometry. Data are normalized to the mean of P25 *Csf1r-cre*^*-*^ *Mertk* ^f/f^ mice and represented as mean ± SEM, n = 6 mice/ group, not significant (n.s.) by Mann-Whitney’s test. **(C)** Representative histograms showing expression of CD68, CD11c and CD86 on microglia from P25 *Csf1r-cre*^*+*^ *Mertk* ^f/f^ and *Csf1r-cre*^*-*^ *Mertk* ^f/f^ mice are shown, next to corresponding quantification of mean fluorescence intensity (MFI) represented as mean ± SEM. N = 6 mice/ group, not significant (n.s.) by Mann-Whitney’s test. **(D)** Schematic of eye transversal sections indicating the intermediate region of the retina wherein ONL thickness was quantified. Eye transversal sections were stained with hematoxylin and eosin. Representative retinal sections and quantification of ONL thickness in the medial portion of the retina. Mean ± SEM of 10 measurements/ mouse, n = 3 mice/ genotype. Not significant (n.s.) *vs. Csf1r-cre*^*-*^ *Mertk* ^f/f^ by Mann-Whitney’s test. Scale bars = 200 μm (whole section) and 10 μm (inset).

### Pharmacological inhibition of JAK kinase activity significantly inhibits photoreceptor cell death and preserves ONL thickness

The observation that a broad-spectrum type I and type II IFN and IL-6 response characterizes the incipient RPE inflammation led us to test whether inhibition of JAK-STAT signaling, a key effector of the type I and II IFN and IL-6 response, could at least partially prevent the PR degeneration in *Mertk* ^-/-V1^ mice. Ruxolitinib is an FDA-approved, selective JAK1/2 inhibitor ^30,31^. Ruxolitinib is known to preferentially partition into milk, reaching a milk:plasma mean concentration ratio of 13.4:1 (drugs.com). To initiate ruxolitinib treatment, ruxolitinib (2g/kg)-containing chow were fed to *Mertk* ^-/-V1^ or B6 WT dams starting at day 10 after parturition so that it is passed on to suckling pups. Weaned pups were continued on ruxolitinib chow until either P25 or P30 (**Fig. 8A**). As controls, nursing dams and their weaned pups were fed standard diet. Following this regimen of ruxolitinib treatment and control, we tested target inhibition in the pups (**Fig. 8B**). Pups were euthanized at P25, mRNA was isolated from RPE, and expression of selected genes was measured by qPCR. These results demonstrate that 15 days on ruxolitinib significantly decreased JAK1/2 driven expression of inflammatory genes such as *Ifit1, Tgtp1* and *Gm4951* in the RPE of the pups, by comparison with no treatment (**Fig. 8B**). Having confirmed target validation, we performed H&E staining on retinal sections at P30 and quantitated the thickness of the ONL in WT B6 and *Mertk* ^-/-V1^ mice with or without ruxolitinib treatment (**Fig. 8C**). While ONL thickness was reduced ∼35% in *Mertk* ^-/-V1^ mice without ruxolitinib treatment, by comparison to untreated B6 WT mice, ONL thickness was reduced only by ∼15% in *Mertk* ^-/-V1^ mice treated with ruxolitinib, by comparison to untreated B6 WT mice (**Fig. 8C**). Thus, ruxolitinib treatment resulted in a statististically significant, ∼2-fold less degeneration of the ONL in *Mertk* ^-/-V1^ mice. As an alternate method for quantifying PR loss, we also counted rows of PR nuclei of the ONL. B6 WT mice had 10.67 ± 0.13 rows of ONL nuclei, while standard chow-treated *Mertk* ^-/-V1^ mice had 6.89 ± 0.06 rows of ONL nuclei *i*.*e*. ∼40% reduction in the number of ONL nuclei. By contrast, ruxolitinib-treated *Mertk* ^-/-V1^ mice had 8.33 ± 0.38 rows of ONL nuclei *i*.*e*. ∼20% reduction in the number of ONL nuclei (**Fig. 8C**). Again, ruxolitinib-treated *Mertk* ^-/-V1^ mice exhibited ∼2-fold less degeneration. Collectively, both forms of quantification demonstrate that the dose (2g/kg) and regimen (from P10 to P30) of ruxolitinib used significantly lessened the severity of the *Mertk* ^-/-V1^ retinal degeneration, albeit PR loss was not entirely prevented.

**Fig. 8.**
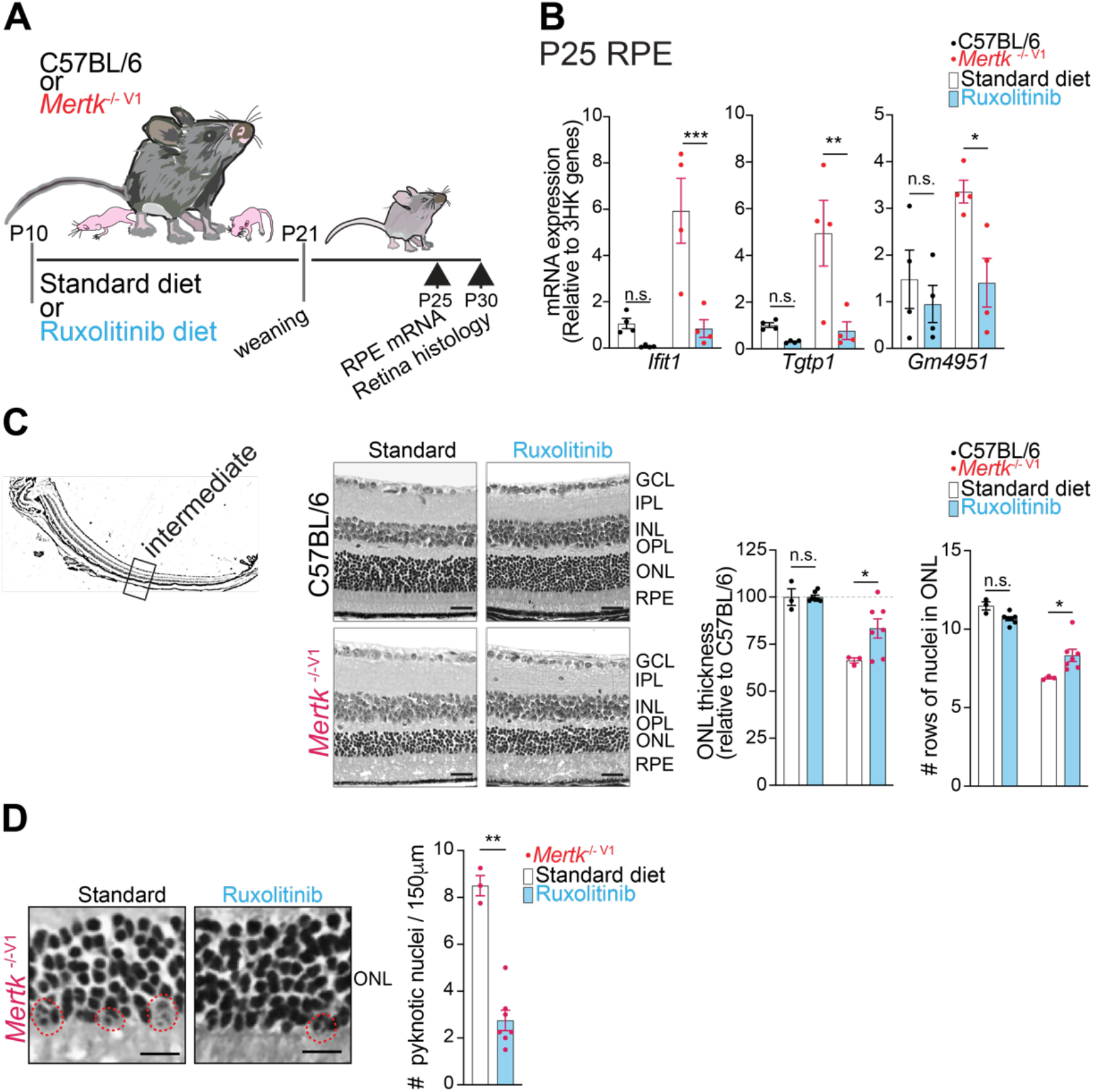
The selective JAK1/2 inhibitor ruxolitinib significantly prevents early-onset PR death. **(A)** Diagram showing experimental design. C57BL/6 WT and *Mertk* ^-/-V1^ dams were fed a standard or ruxolitinib-containing diet (2g ruxolitinib/ kg of chow) from the time pups were at postnatal day (P) 10 until weaning at P21. Weanlings continued to receive standard or ruxolitinib-containing diet until the indicated times of sample collection. RPE samples were collected at P25 for mRNA expression of target genes. Eyes were collected at P30 to study retinal histology. **(B)** qPCR of inflammation-associated genes was performed following collection of RPE at P25. Data are represented as mean ± SEM n = 4 mice/ condition. Not significant (n.s.), *p < 0.05, **p < 0.01, ***p < 0.001 by two-way ANOVA. **(C)** Schematic of eye transversal sections indicating the intermediate region of the retina wherein ONL thickness was quantified. Representative eye sections from C57BL/6 WT and *Mertk* ^-/-V1^ mice collected at P30 and stained with hematoxylin and eosin after treatment with standard or ruxolitinib or standard diet, and quantification of ONL thickness and number of rows of nuclei within ONL are shown. Mean ± SEM n = 3-7 mice/ condition. Not significant (n.s.), *p < 0.05 by two-way ANOVA. Scale bars = 20 μm. **(D)** Representative images and quantification of the number of pyknotic nuclei in ONL of P30 *Mertk* ^-/-V1^ mice treated with ruxolitinib or standard diet. Data are represented as mean ± SEM n = 3-7 mice/ condition. **p < 0.01 by Mann-Whitney’s test. Scale bars = 10 μm.

We also quantitated the pyknotic nuclei within the ONL of *Mertk* ^-/-V1^ mice fed on standard or ruxolitinib chow as a measure of dying PR (**Fig. 8D**). These results showed that there was an ∼4-fold reduction in the number of pyknotic nuclei with ONL of *Mertk* ^-/-V1^ mice with ruxolitinib treatment, by comparison to untreated *Mertk* ^-/-V1^ mice (**Fig. 8D**). Thus, subduing RPE inflammation by inhibition of JAK1/2 activity at least partially but significantly reduces the severity of the aggressive, early-onset PR degeneration phenotype observed in *Mertk* ^-/-V1^ mice.

## Discussion

### Severe, early-onset PR degeneration due to MERTK deficiency is associated with RPE inflammation, microglial activation and monocyte infiltration into the retina

RP associated with *MERTK* mutations is a severe, early-onset form of this disease. It has been speculated that RP with *MERTK* mutations is associated with inflammation – nonetheless, the role of inflammation in disease etiology was less clear. Here we show that *Mertk* and *Tyro3* function redundantly as negative regulators of inflammation in the mouse RPE. Mice lacking WT expression of *Mertk* and hypomorphic expression/loss of *Tyro3* displayed a broad-spectrum RPE inflammatory response including type I and II IFN response, IL-6 signaling pathway gene expression and TNF-α response gene expression. Microglial activation was also a characteristic of retinal inflammation in *Mertk* ^-/-V1^ mice, and phenocopied in *Mertk* ^-/-V2^ *Tyro3* ^-/-V2^ mice. Both these mouse models were also associated with monocyte infiltration into the retina. This cascading inflammation culminated in severe, early-onset retinal degeneration. By contrast, mouse models of defective phagocytosis that were not associated with RPE inflammation, such as *Mertk* ^-/-V2^ mice or the phenotype of the *Itgb5* ^-/-14^ mice, presented with an age-dependent onset and progressive changes such as thinning of the peripheral retina. Thus, RPE inflammatory response may be a crucial feature of at least a subset of *MERTK* mutation-associated RP.

### Inflammation is not a consequence of failed phagocytosis

Inflammation, heretofore, has been mostly considered a secondary consequence of failed phagocytosis, accumulation of damaged and degraded POS and the death of PRs. We showed recently that neural retina-resident microglia are already activated in RCS rat at an age prior to detectable PR death and debris buildup ^26^. Here, our characterization of the inflammatory response in the RPE itself revealed that not only does RPE inflammation precede retinal degeneration, but also that the inception of RPE inflammation happens before eye opening in mouse pups, a time when phagocytosis is expected to be minimal ^26^. If RPE inflammation were to be only a consequence of a failure to clear POS debris and accumulation of dead PR, such a process can be expected to lag the primary defect in phagocytosis. A gradual process of build-up of POS and dead PRs would progressively culminate in RPE inflammation. Instead, we observe the opposite, *i*.*e*. histological changes in the retina trail the initial RPE inflammatory response. We also observe that inflammation and PR degeneration are not merely obligatory consequences of defective phagocytosis. This notion is supported by the finding that loss of function of the phagocytic receptor *Itgb5* did not phenocopy the inflammation observed in *Mertk* ^-/-V1^ or *Mertk* ^-/-V2^ *Tyro3* ^-/-V2^ mice. While only minimal RPE inflammation was seen in *Mertk* ^-/-V2^ mice, RPE from these mice were deficient in phagocytosis. *Mertk* ^-/-V2^ mice had demonstrable phagocytosis defects in their macrophages as well, consistent with the phagocytic function of MERTK ^27^. Neither did we observe increased inflammation in the *Tyro3* ^-/-V1^ RPE, although TYRO3 is reportedly involved in POS phagocytosis ^17^. Interestingly, *Itgb5* ^-/-^, *Mertk* ^-/-V2^ or *Tyro3* ^-/-V1^ mice do not develop severe, early-onset PR degeneration ^14,27^. Thus, impaired peak phagocytosis is not intrinsically associated with an enhanced inflammatory response gene expression and PR degeneration. Consistent with this idea, a previous study had demonstrated that the ectopic expression of another phagocytic receptor BAI-1 in *Mertk* ^-/-v1^ mice failed to prevent or reduce PR degeneration ^32^. MERTK and TYRO3 not only participate in phagocytosis, but also function in the negative regulation of inflammation ^33^. These receptor tyrosine kinases (RTKs) have a Grb2-binding motif that is functionally essential for their phagocytic function ^34^, but also a distinct immunoreceptor tyrosine-based inhibitory (ITIM) motif ^33^. Our results indicate that RPE homeostasis requires the redundant function of these RTKs for the negative regulation of inflammation. It will be interesting to ascertain the individual roles of mutating these motifs in MERTK and TYRO3 in determining the severe, early-onset PR degeneration phenotype associated with the loss of *Mertk* and the hypomorphic expression/loss of *Tyro3*.

### RPE *versus* microglial inflammation as the nucleating event for PR degeneration

Although microglia (and even other myeloid cells) are involved in a cascading inflammatory response within the retina leading to PR degeneration, we and others have demonstrated that the severe, early-onset PR degeneration phenotype in *Mertk* ^-/-V1^ and *Mertk* ^-/-V2^ *Tyro3* ^-/-V2^ mice requires both the loss of *Mertk* as well as the concomitant hypomorphic expression/simultaneous genetic ablation of *Tyro3* ^17,27^. While *Mertk* is expressed in microglia and widely in myeloid cells ^33^, *Tyro3* expression has only been described in very specific myeloid populations such as CD11c^+^ PDL2^+^ dendritic cells involved in the inhibition of type 2 immunity, such as in asthma or allergic diseases ^35^. Therefore, we hypothesized that rather than microglia or myeloid cells, inflammation associated with severe, early-onset PR degeneration likely nucleates in a tissue that expresses both *Mertk* and *Tyro3*. This made us investigate RPE inflammation. Two pieces of independent evidence support the RPE as one of the initial sites of an inflammatory response culminating in severe, early-onset PR degeneration. First, RPE inflammation was detected at P10, before microglial activation. Second, the *Csf1r*-cre-dependent genetic ablation of *Mertk*, which ablates this RTK in microglia and in myeloid cells, did not result in an inflammatory response and retinal degeneration. However, it is evident that microglial activation is a part of the inflammatory response that occurs as a reaction to microglia-extrinsic inflammation. In our study on MERTK-deficient RCS rats, inhibition of retinal microglial activation with tamoxifen delayed retinal degeneration, despite the fact that the efficacy of tamoxifen was moderate ^26^. A head-to-head comparison of targeting RPE nucleated inflammation *via* ruxolitinib *versus* microglial activation by tamoxifen in RCS rats or in *Mertk*^-/-V1^ and *Mertk* ^-/-V2^ *Tyro3* ^-/-V2^ mice or testing a combination approach has not yet been done. In other models of retinal degeneration such as acute light damage and the rd10 model of RP wherein rods due to lack of PDE6 degenerate between P20 to P25 ^36^, mice treated with tamoxifen had improved light responses compared to untreated mice illustrating the common involvement of microglia in retinal disease with different cause and etiology ^37^. Whether RPE inflammation is also shared among different forms of retinal degeneration, and if so whether targeting RPE inflammation would be more effective than targeting microglia, is as yet unexplored.

The primordial trigger of inflammation, however, remains undiscovered. The visual cycle is intrinsically associated with inflammation. Following photobleaching, rod cells return to their basal state as all-*trans*-retinal is removed from disk membranes, cGMP is re-synthesized and rhodopsin is regenerated from 11-*cis*-retinal and opsin ^38-40^. The RPE plays a role in uptake of all-*trans*-retinol transported out of POS and its isomerization into 11-*cis*-retinol followed by its oxidization to 11-*cis*-retinal. Mutations in genes encoding transporters such as *ABCA4* involved in the elimination of all-*trans*-retinal from photoreceptors are known to cause macular degeneration, rod-cone dystrophies, or RP ^41,42^. All-*trans*-retinal has also been reported to activate the NLRP3 inflammasome and cause RPE death ^43^. Thus, RPE cells can be expected to require anti-inflammatory mechanisms for the uptake and recycling of inflammatory products of the visual cycle eliminated by the photoreceptors, such as all-*trans*-retinol. Spontaneous electrical activity in the retina has been reported before eye opening ^44^ indicating that phototoxic damage and the requirement to keep inflammation at bay may be expected to be required quite early in development. Still, we cannot rule out the loss of negative regulation of inflammation in other cell types expressing both *Mertk* and *Tyro3*, such as Müller glia, as the genesis that spills over to the RPE.

### Inflammation as the target of therapy in severe, early-onset RP due to MERTK mutation

The molecular characteristic of RPE inflammation afflicting *Mertk* ^-/-V1^ or *Mertk* ^-/-V2^ *Tyro3* ^-/-V2^ mice was enhanced type I and II IFN, IL-6 and TNF-α signaling. The pathogenic role of TNF-α in *Mertk* loss of function in murine RP was previously ruled out ^23^. Both type I and II IFN and IL-6 pathways are dependent on the JAK kinases, JAK1/2. We tested whether JAK1/2 inhibition would ameliorate the severe, early-onset PR degeneration. Indeed, a treatment regimen with ruxolitinib from P10 to P30 significantly reduced PR death and the shrinking of the ONL. Whether administration of ruxolitinib earlier would provide improved benefit and more protection against PR degeneration is yet to be determined. Collectively, our results demonstrate that in the absence of *Mertk* and deficits of *Tyro3*, a nascent JAK1/2-STAT-dependent inflammatory response pathway within RPE leads to a panoply of cytokines, chemokines, activation of microglia, microglial proliferation, microglial migration and eventually, a runaway cascade of inflammatory response involving the infiltration of monocytes, to culminate in severe, early-onset PR degeneration. Whether other modifiers influence an incipient RPE inflammatory response in RCS rat or in human RP patients remains to be investigated. Furthermore, the nature of the inflammatory response and whether it is amenable to therapeutic targeting as in *Mertk* ^-/-V1^ mice remain unknown. The presence of *MERTK* mutations that do not result in severe, early-onset RP appears to indicate that modifiers determine the severity and age of onset of *MERTK* mutation-associated RP. If indeed modifiers of *MERTK-*associated RP phenotype govern an inflammatory component responsible for severe, early-onset retinal degeneration, inhibiting this inflammatory axis may provide a promising, tractable avenue for therapeutic benefits.

## Materials and Methods

### Experimental Design

This study was designed to identify the pathological drivers of early-onset PR degeneration in mice with genetic ablation of *Mertk* and *Tyro3*. We employed transcriptional, histological, immunological, and functional analysis of RPE and retina. We used mice at different ages, ranging from postnatal day (P) 10 to 12-month-old mice. We tested the efficacy of the FDA-approved JAK1/2 inhibitor ruxolitinib in the prevention of early-onset retinal degeneration in mice with loss of *Mertk* and hypomorphic expression of *Tyro3*. Experiments were independently replicated and for each experiment an n > 4 for each group, unless otherwise indicated in the figure legends, was used.

### Animals

Animals were bred and maintained under a strict 12-hour light cycle and fed with standard chow diet in a specific pathogen-free facility at Yale University. All animal experiments were performed in accordance with regulatory guidelines and standards set by the Institutional Animal Care and Use Committee of Yale University and Fordham University. C57BL/6J mice were purchased from Jackson laboratories (strain #:000664) and subsequently bred and housed at Yale University. *Mertk* ^-/-V1^ mice have been described previously ^22,45,46^. The generation of *Mertk* ^-/-V2^ and *Mertk* ^-/-V2^ *Tyro3* ^-/-V2^ mice is described in Akalu *et al*., 2022 ^22^. *Tyro3* ^-/-V1^ mice have also been described previously ^46^. *Mertk* ^-/-V1^, *Mertk* ^-/-V2^ and *Mertk* ^-/-V2^ *Tyro3* ^-/-V2^ mice, as well as *Csf1r-cre Mertk* ^*f*/f^ mice (bearing conditional ablation of *Mertk* in myeloid cells) ^47^ were bred and housed at Yale University. Previously described ^14^ *Itgb5*^-/-^ mice and strain-matched 129T2/SvEmsJ WT controls (Jackson laboratories strain#:002065) were bred and housed at Fordham University. Genotyping of selected animals of the various aforementioned *Mertk, Tyro3 or Itgb5* genotypes in our colonies showed them all to be negative for the *rd8* mutation. Genome wide SNP analysis for *Mertk* ^-/-V2^ mice determined that ∼84.37% of the genome in *Mertk* ^-/-V2^ mice corresponded to C57BL/6J and ∼15.62% to C57BL/6NJ ^22^.

### Total RNA isolation and sequencing analysis

RNA was collected from P25 mice using a previously validated method ^48^. Briefly, after euthanasia, mice were enucleated and the posterior eyecup was incubated on ice in 200 μl of RNAprotect (Qiagen) for 1 hour. RPE containing tubes were agitated for 10 minutes to dislodge any RPE cells attached to the posterior eyecup and centrifuged for 5 minutes at 685g. The RPE pellet was then subjected to total RNA extraction using RNeasy Mini kit (Qiagen) following manufacturer’s instructions.

RNA libraries were prepared at the Yale Keck Biotechnology Resource Laboratory from three to six biological replicates per condition. Samples were sequenced using 150 bp paired-end reading on a NovaSeq 6000 instrument (Illumina). RNA-sequencing datasets from P25 RPE obtained from C57BL/6, *Mertk*^-/-V1^, and *Mertk*^-/-V2^ mice are available through the Gene Expression Omnibus (GEO) under accession ID: GSE205070 ^22^. Datasets corresponding to P25 RPE from *Mertk*^-/-V2^ *Tyro3*^-/-V2^, *Tyro3*^-/-V1^, *Igtb5*^-/-^ and WT^129^ mice are available through GEO under accession ID: GSE214005. The raw reads were then subjected to trimming by btrim ^49^ to remove sequencing adaptors and low-quality regions. Next, reads were mapped to the mouse genome (GRCm38) using STAR ^50^. Finally, the DEseq2 ^51^ package was run to identify differentially expressed genes according to p-values adjusted for multiple comparisons. Genes with p-adjusted values less than 0.05 and log_2_fold change ≤-1.25 or ≥1.25 were considered differentially expressed. These data were subsequently represented in a heatmap generated by Qlucore Omics Explorer. Volcano plots were generated using GraphPad Prism (GraphPad Software Inc.). We also subjected normalized counts of all genes from various samples to Gene set enrichment analysis ^52,53^. The hallmark gene set database was used to determine which pathways are enriched in our samples. Resulting pathways that had high normalized enrichment scores and FDR Q value <0.05 were reported.

### Quantitative PCR analysis

Reverse transcription of RNA was performed utilizing iScript cDNA Synthesis Kit (Bio-Rad). Using KAPA SYBR Fast qPCR Kit (Kapa Biosystems), we amplified cDNA fragments and proceeded with qPCR reactions on CFX96 Thermal Cycler Real Time System (Bio-Rad). The reactions were normalized to 3 housekeeping genes (*Gapdh, Hprt* and *Rn18s*) and specificity of the amplified products was verified by looking at the dissociation curves. All oligonucleotides for qPCR were either purchased from Sigma-Aldrich or produced at the Yale University Keck Oligonucleotide Synthesis Facility (see sequences in Table S1).

### Luminex cytokine assay

Eyes from P25 mice were obtained. Neural retinas were subsequently separated from posterior eyecups and both tissues were incubated on ice in RIPA buffer with protease inhibitor cocktail, EDTA-free (Sigma-Aldrich, Inc.) for up to 1 hour. Next, posterior eyecups were removed and the dislodged RPE cells were combined with corresponding neural retinas and sonicated for 20 seconds on ice. After 10 minutes at 14,000 rpm in a refrigerated centrifuge, supernatants were transferred to new tubes. Undiluted supernatants from n = 5 mice/genotype were then tested for expression of 44 different cytokines and chemokines on the Mouse Cytokine 44-Plex Discovery Array (Eve Technologies). RPE lysates were also tested independently for cytokine expression using the same platform.

### Histological analysis

After sacrifice by carbon dioxide inhalation, eyecups were immediately collected, incubated overnight in eye fixative (ServiceBio). Hematoxylin and eosin staining was performed by iHisto Inc. Samples were processed, embedded in paraffin, and sectioned at 4 μm. Paraffin sections were then deparaffinized and hydrated using the following steps: 15 minutes in xylene twice; 5, 5, and 5 minutes in 100%, 100%, and 75% ethanol, respectively; and 5 minutes in 1x PBS at room temperature repeated three times. After deparaffinization, 4-μm-sectioned samples were placed on glass slides and stained with hematoxylin and eosin. Whole slide scanning (20x) was performed on an EasyScan Infinity (Motic). Outer nuclear layer (ONL) thickness was analyzed in ImageJ (NIH). Quantification of ONL thickness was performed in the central, medial or peripheral portions of the retina, as indicated in the corresponding figure legends. The areas analyzed were defined by distance from the optic disk. 10-15 measurements of ONL thickness were done per mouse. PR length, the combined length of inner and outer PR segments, was quantified and averaged from 10 measurements from 2-3 eye sections per mouse using ImageJ (NIH).

### Immunofluorescence

For IBA1 staining, 4 μm paraffin sections were deparaffinized at 65°C for 20 minutes, and incubated in xylenes for 5 minutes twice. Sections were subsequently incubated in 100% ethanol (2x 5 minutes), 95% ethanol (2x 3 minutes), 70% ethanol (3 minutes) and 5 minutes in PBS at room temperature. Epitope unmasking was performed using citrate buffer (10 mM citric acid, 0.05% Tween-20, pH 6) for 10 minutes in the microwave (10% total power). Sections were then blocked in PBS, 1% BSA with 0.01% Triton-X100 for 30 minutes at room temperature and incubated overnight at 4°C with primary antibody anti-IBA1 (1:400, Wako Chemicals USA). After washing, samples were incubated with Donkey anti-rabbit Alexa Fluor 488-conjugated secondary antibody (1:500, Thermo Fisher Scientific) for 1 hour at room temperature and mounted on slides using DAPI-containing mounting media (Vector Laboratories). Image acquisition was performed using a Leica Stellaris 8 confocal microscope using an oil-immersion 40X objective. The layers of the outer retina were delineated using DAPI, and number of IBA1+ cells were counted within each layer (INL, OPL, ONL). Images were analyzed using ImageJ software (NIH).

### Electroretinography (ERG) recordings

All experimental animals were adapted in a dark room for 12 hours prior to recordings. Animals were anesthetized under dim red illumination using a 100 mg/kg Ketamine and 10 mg/kg xylazine cocktail injected intraperitoneally and pupils were dilated by application of a 0.5% tropicamide eye drop (Sandoz) at least 15 minutes before recordings. The cornea was intermittently irrigated with balanced salt solution to maintain the baseline recording and prevent keratopathy. Scotopic electroretinograms were acquired with UTAS ERG System with a BigShot Ganzfeld Stimulator (LKC Technologies, Inc.). A needle reference electrode was placed under the skin of the back of the head, a ground electrode was attached subcutaneously to the tail, and a lens electrode was placed in contact with the central cornea. The scotopic response was recorded for different luminances (i.e., log -2.1, -0.6, 0.4 and 1.4 cd.s/m^2^) using EMWin software, following manufacturer’s instructions (LKC Technologies, Inc.). The a-wave was measured as the difference in amplitude between baseline recording and the trough of the negative deflection, and the b-wave amplitude was measured from the trough of the a-wave to the peak of the ERG.

### Whole-mount RPE preparation, labeling, and *in situ* phagosome quantification

Quantification of POS phagosomes in whole-mount RPE *in situ* was performed as described previously ^54^. Tissues were collected immediately post-mortem from 3-month-old mice sacrificed at 1 and 7 hours after light onset (n = 5 of each genotype). Posterior eyecups were dissected and fixed for 30 minutes in 4% paraformaldehyde in PBS. Radial cuts were made to flatten tissues before labeling with rhodopsin monoclonal antibody B6-30 (Abcam) ^55^ and AlexaFluor 488-conjugated donkey anti-mouse secondary antibody. Counterstains labeling nuclei with DAPI and F-actin with AlexaFluor594-conjugated phalloidin (Thermofisher) served to indicate cell orientation and integrity. A Leica TSP8 confocal system was used to acquire x-y image stacks at 0.25 μm intervals to cover the thickness of the entire RPE depth from the basal aspect of nuclei to apical microvilli of the RPE. 13-16 fields were imaged for each specimen with 186.5 μm x 186.5 μm field size. Fields were spaced along the dorsoventral and nasotemporal retinal meridians. Maximal projections of z-stacks of equal thickness were used to quantify POS phagosomes defined by rhodopsin-positive structures of ≥0.5 μm or 0.9 μm diameter using ImageJ.

### Flow cytometry staining and acquisition

The method used to isolate and immunophenotype the retina was previously described ^56^. Briefly, retinas were digested for 1 hour at 37°C with agitation in RPMI (Gibco) containing 1.5mg/ml of Collagenase A (Roche) and 0.4 mg/ml of DNaseI (Sigma-Aldrich). For adult P42 mice, retinas were collected after intracardiac perfusion with saline solution. Single cell suspensions of tissues were then stained in PBS with Zombie viability dye (BioLegend) for 10 minutes. Cells were then incubated in 2% FCS/PBS solution containing anti-mouse CD16/32 antibody (BioLegend clone 93) for 15 minutes and subsequently stained with a combination of fluorophore-conjugated primary antibodies against mouse CD45 (BioLegend clone 30-F11), CD11b (BioLegend clone M1/70), CD64 (BioLegend clone X54-5/7.1), Ly6C (BioLegend clone HK1.4), Ly6G (BioLegend clone 1A8), CD68 (BioLegend clone FA-11), Galectin-3 (BioLegend clone M3/38), CD11c (BioLegend clone N418), and CD86 (BioLegend clone GL-1) at 4°C for 25 minutes. After staining, cells were washed and fixed in 1% paraformaldehyde in PBS. Data were acquired with BD LSRII flow cytometer using BD FACSDiva software (BD Biosciences). Finally, raw data were analyzed using FlowJo software (Tree Star Inc).

### Ruxolitinib treatment

Ruxolitinib is an FDA-approved, selective JAK1/2 inhibitor used to treat intermediate or high-risk myelofibrosis ^30^. Ruxolitinib was administered to C57BL/6 or *Mertk* ^-/-V1^ mice, by providing ruxolitinib-supplemented chow (2 g/kg, Research diets) *ad libitum* to the nursing dams starting at P10 (pups received the drug indirectly through breast milk), and after that directly to the experimental mice from weaning (P21) until sacrifice. Control littermates were manipulated identically but received standard chow (Envigo) throughout the duration of the experiments. RPE samples were collected at P25 for mRNA studies or at P35 for histological evaluation.

### Statistical analysis

All statistical analyses were done using GraphPad Prism (GraphPad Software Inc.). All data are shown as mean ± S.E.M. and each data point represents a unique animal. Statistical differences between experimental groups were determined by employing non-parametric tests— namely, one and two-tailed Mann-Whitney test, two-tailed Student’s t-test, one-way and two-way ANOVAs.

## Acknowledgments

The authors would like to acknowledge the members of the Rothlin-Ghosh laboratory for scientific discussions relating to the preparation of this manuscript. Flow cytometry experiments were done at the Yale Flow Cytometry Facility.

## Author contributions

Conceptualization: MEM, YTA, FM, GG, SCF, CVR, SG

Methodology: MEM, YTA, FM, GG, EA, YK, BPH

Investigation: MEM, YTA, FM, GG, EA

Visualization: MEM, YTA, FM, GG

Supervision: BPH, SCF, CVR, SG

Writing—original draft: MEM, YTA, SCF, CVR, SG

Funding acquisition: SCF, CVR, SG

## Data and materials availability

The accession numbers for the RPE bulk RNA sequencing data reported in this paper are GEO: GSE205070 ^22^ and GSE214005. Further information and requests for resources and reagents should be directed to Principal Investigators, Carla V. Rothlin and Sourav Ghosh.

