## Supplementary materials for "Inflammation of the retinal pigment epithelium drives early-onset photoreceptor degeneration in *Mertk*-associated retinitis pigmentosa"

**This PDF file includes:**

Figs. S1 to S5
Table S1

**Fig. S1. Preserved ERG function in 12-months old *Mertk* $^{-/-V2}$ and *Tyro3* $^{-/-V1}$ mice.**

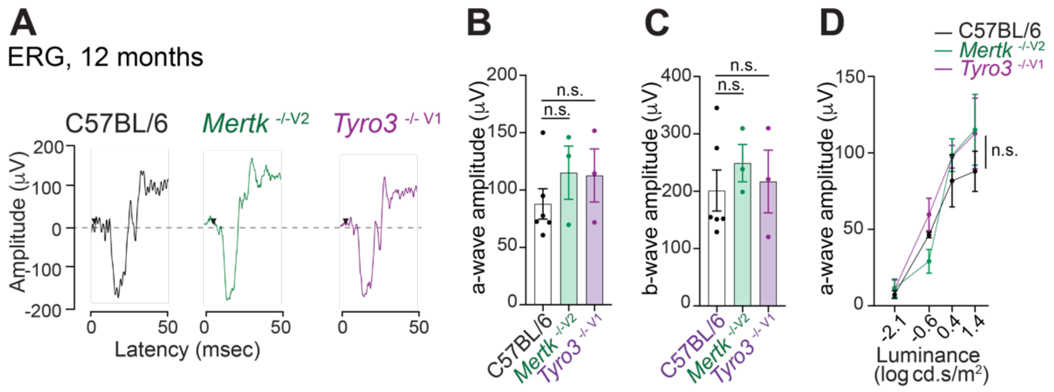


**(A)** Representative scotopic ERG traces from 12-month-old C57BL/6 WT, *Mertk* $^{-/-V2}$ and *Tyro3* $^{-/-V1}$ mice are shown at the highest luminance tested (25 cd.s.m$^{-2}$).

**(B)** Quantification of a-wave amplitude at 25 cd.s.m$^{-2}$. Data are represented as mean ± SEM, n = 3-6 mice/ genotype. Not significant (n.s.) *vs.* C57BL/6 WT by one-way ANOVA.

**(C)** Quantification of b-wave amplitude at 25 cd.s.m$^{-2}$. Data are represented as mean ± SEM, n = 3-6 mice/ genotype. Not significant (n.s.) *vs.* C57BL/6 WT by one-way ANOVA, followed by Dunnet's test.

**(D)** Quantification of a-wave amplitudes at increasing luminance. Data are represented as mean ± SEM, n = 3-6 mice/ genotype. Not significant (n.s.) *vs.* C57BL/6 WT by two-way ANOVA.

**Fig. S2. Proliferation and activation of retinal microglia and monocyte infiltration are increased in retinas from *Mertk* $^{-/-V1}$ and *Mertk* $^{-/-V2}$ *Tyro3* $^{-/-V2}$ mice at post-natal day 42.**

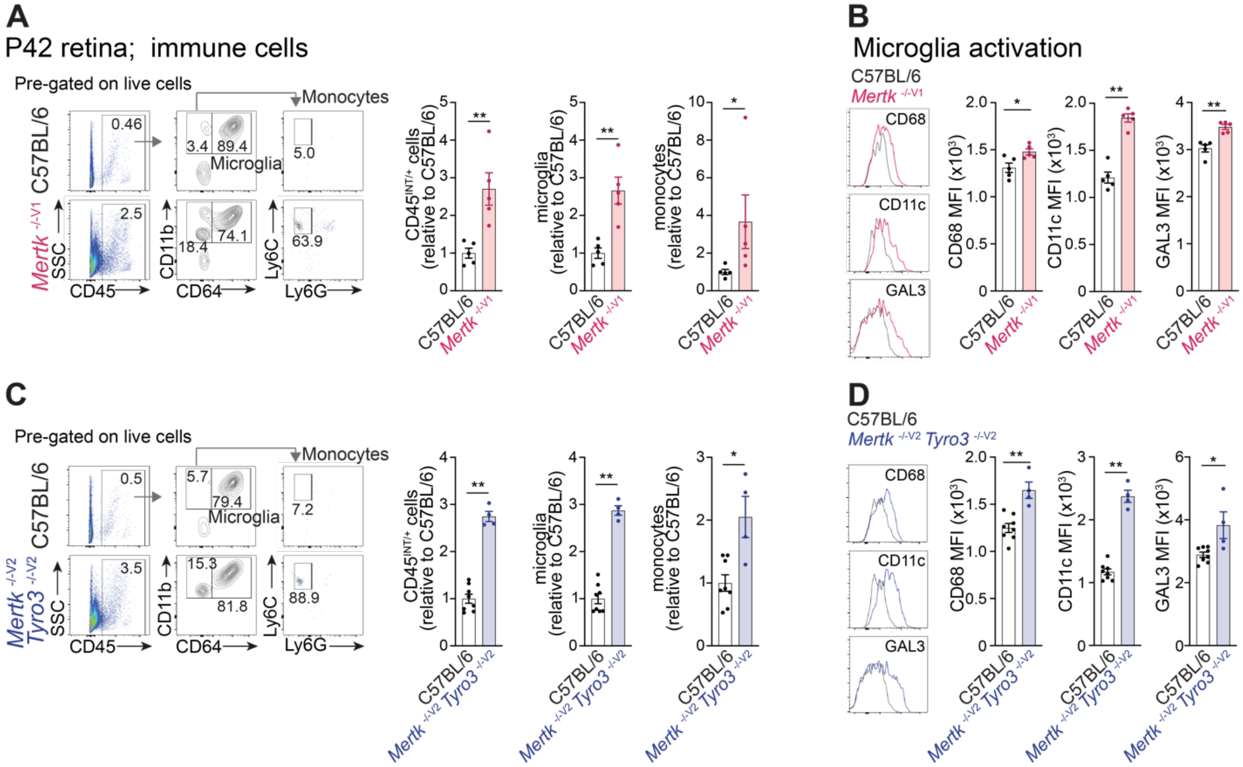


**(A)** Retinas from P42 C57BL/6 WT and *Mertk* $^{-/-V1}$ were dissociated, and the type, number and activation of immune cells were determined by flow cytometry. The number of CD45$^{INT/+}$ cells, microglia (CD45$^{INT/+}$ CD11b$^{+}$ CD64$^{+}$ cells) and infiltrating monocytes (CD45$^{+}$ CD11b$^{+}$ CD64$^{-}$ Ly6C$^{+}$ Ly6G$^{-}$ cells) within retinas is normalized to the mean of C57BL/6 WT and represented as mean ± SEM, n = 5 mice/ group. *p < 0.05 and **p < 0.01, by Mann-Whitney's test.

**(B)** Representative histograms showing expression of CD68, CD11c and GAL3 on microglia are shown, next to corresponding quantification of mean fluorescence intensity (MFI) represented as mean ± SEM, n = 5 mice/ group. *p < 0.05, and **p < 0.01 by Mann-Whitney's test.

**(C)** Retinas from P42 C57BL/6 WT and *Mertk* $^{-/-V2}$ *Tyro3* $^{-/-V2}$ were dissociated, and the type, number and activation of immune cells were determined by flow cytometry. The number of CD45$^{INT/+}$ cells, microglia (CD45$^{INT/+}$ CD11b$^{+}$ CD64$^{+}$ cells) and infiltrating monocytes (CD45$^{+}$ CD11b$^{+}$ CD64$^{-}$ Ly6C$^{+}$ Ly6G$^{-}$ cells) within retinas is normalized to the mean of

C57BL/6 WT mice and represented as mean ± SEM, n = 4 - 8 mice/ group, *p < 0.05, and **p < 0.01 by Mann-Whitney's test.

**(D)** Representative histograms showing expression of CD68, CD11c and GAL3 on microglia are shown, next to corresponding quantification of mean fluorescence intensity (MFI) represented as mean ± SEM, n = 4 - 8 mice/ group, *p < 0.05, and **p < 0.01 by Mann-Whitney's test.

**Fig. S3. RPE morphology is preserved in *Mertk* $^{-/-V2}$ mice.**

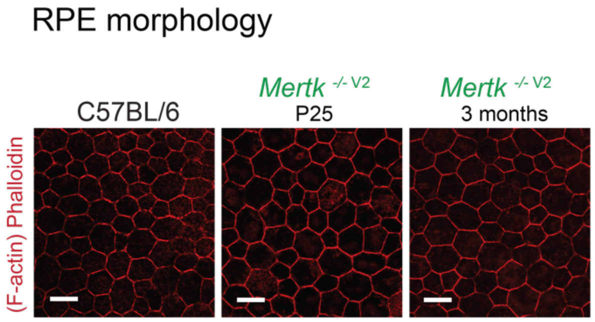


Representative images of phalloidin (F-actin) staining in whole-mount RPE in P25 and 3-month-old C57BL/6 WT and *Mertk* $^{-/-V2}$ mice showing preserved RPE morphology in *Mertk* $^{-/-V2}$ mice. Scale bars = 20 μm.

**Fig. S4. Deficiency of phagocytic receptor subunit *Itgb5* does not lead to early-onset RPE inflammation.**

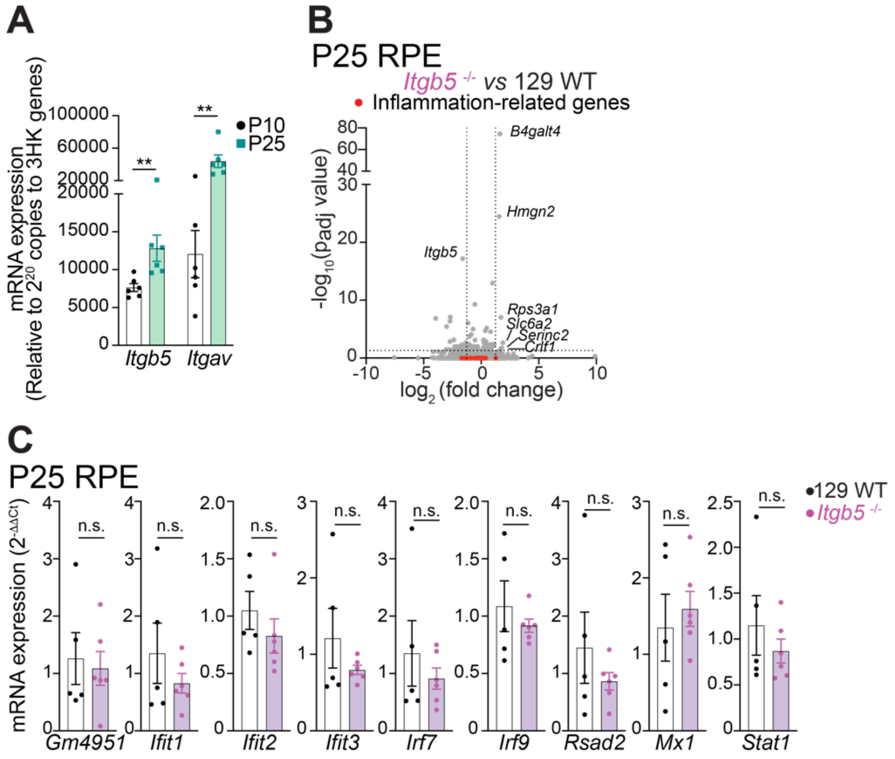


**(A)** qPCR measurement of amounts of *Itgb5* and *Itgav* mRNA expression in RPE cells collected from C57BL/6 WT mice at P10 and P25 (mean ± SEM, n = 5 - 6 mice/ group, $^{**}p < 0.01$ by Mann-Whitney's test).

**(B)** Volcano plot comparing gene expression at P25 in *Itgb5* $^{-/-}$ RPE relative to 129 WT RPE by RNA sequencing (n = 3 samples/ genotype, each sample was comprised of pooled tissues from 2 animals). Dashed lines indicate thresholds for significance (FC>2 or FC<-2 and padj<0.05). Genes indicated in red are associated with inflammation and they have been shown to be upregulated in *Mertk* $^{-/-V1}$ and/or *Mertk* $^{-/-V2}$ *Tyro3* $^{-/-V2}$ RPEs (Figure 3).

**(C)** Assessment of interferon-inducible genes at P25 in *Itgb5* $^{-/-}$ RPE relative to 129 WT RPE by qPCR (mean ± SEM, n = 5 - 6 mice/ group, not significant (n.s.) by Mann-Whitney's test).

**Fig. S5. Efficiency of *Mertk* ablation in *Csf1r-cre*$^{+}$ *Mertk* $^{f/f}$ mice.**

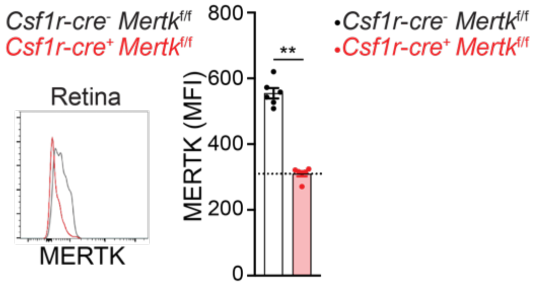


Retinal tissue was isolated from P25 *Csf1r-cre*$^{+}$ *Mertk* $^{f/f}$ and *Csf1r-cre*$^{-}$ *Mertk* $^{f/f}$ mice and immune cells were assessed by flow cytometry. Representative histogram and quantification of MERTK expression in microglia from *Csf1r-cre*$^{+}$ *Mertk* $^{f/f}$ and *Csf1r-cre*$^{-}$ *Mertk* $^{f/f}$ mice are shown (n = 6 mice/ group, **p < 0.01 by Mann-Whitney's test).

**Table S1. Oligonucleotide sequences used for qPCR**

| *Oligonucleotide* | *Sequence* |
|---|---|
| *Cxcl10* F | CCAAGTGCTGCCGTCATTTTC |
| *Cxcl10* R | GGCTCGCAGGGATGATTTCAA |
| *Ifit1* F | CCCAGAGAACAGCTACCACCTT |
| *Ifit1* R | TTGTGCATCCCCAATGGGTT |
| *Ifit3* F | GCTCAGGCTTACGTTGACAAGG |
| *Ifit3* R | CTTTAGGCGTGTCCATCCTTCC |
| *Iigp1* F | GGGCAGGAGTGGATTTCATTGGA |
| *Iigp1* R | ACCTTCACTGCTACAGGCTCTT |
| *Rsad2* F | GGTGCCTGAATCTAACCAGAAG |
| *Rsad2* R | CCACGCCAACATCCAGAATA |
| *Ccl2* F | TTAAAAACCTGGATCGGAACCAA |
| *Ccl2* R | GCATTAGCTTCAGATTTACGGGT |
| *Ccl3* F | CATATGGAGCTGACACCCCG |
| *Ccl3* R | GAGCAAAGGCTGCTGGTTTC |
| *Gm4951* F | CAAGCCAAGAGCACACCGAG |
| *Gm4951* R | GCTCCGCAGCAGGTTATAGCA |
| *Mx1* F | GTGGTAGTCCCCAGCAATGT |
| *Mx1* R | TGCTGACCTCTGCACTTGAC |
| *Irf7 F* | ACAGGGCGTTTTATCTTGCG |
| *Irf7* R | TCCAAGCTCCCGGCTAAGT |
| *Irf9* F | TTTCTTCAGCTCCGCCCTTGC |
| *Irf9* R | ATGGTCTTGGCTGCATCGTC |
| *Lif* F | AACCAGATCAAGAATCAACTGGC |
| *Lif* R | TGTTAGGCGCACATAGCTTTT |
| *Tnfsf9* F | CCCTGTTTCCCACATTGGCTGC |
| *Tnfsf9* R | CTCCATCTTGGCTGTGCCAGTT |
| *Itgav* F | CCGTGGACTTCTTCGAGCC |

| | |
|---|---|
| *Itgav* R | CTGTTGAATCAAACTCAATGGGC |
| *Itgb5* F | GCTGCTGTCTGCAAGGAGAA |
| *Itgb5* R | AAGCAAGGCAAGCGATGGA |
| *Gapdh* F | TCCCACTCTTCCACCTTCGA |
| *Gapdh* R | AGTTGGGATAGGGCCTCTCTT |
| *Hprt* F | AAGCTTGCTGGTGAAAAGGA |
| *Hprt* R | TTGCGCTCATCTTAGGCTTT |
| *Rn18s* F | GTAACCCGTTGAACCCCATT |
| *Rn18s* R | CCATCCAATCGGTACTAGCG |
